# The genomics and physiology of abiotic stressors associated with global elevation gradients in *Arabidopsis thaliana*

**DOI:** 10.1101/2022.03.22.485410

**Authors:** Diana Gamba, Claire Lorts, Asnake Haile, Seema Sahay, Lua Lopez, Tian Xia, Margarita Takou, Evelyn Kulesza, Dinakaran Elango, Jeffrey Kerby, Mistire Yifru, Collins E. Bulafu, Tigist Wondimu, Katarzyna Glowacka, Jesse R. Lasky

## Abstract

Phenotypic and genetic diversity in *Arabidopsis thaliana* may be associated with adaptation along its wide elevational range. We took a multi-regional view of elevational adaptation and in a diverse panel of ecotypes measured plant responses to high elevation stressors: low partial CO_2_ pressure, high light, and night freezing. We conducted genome-wide association studies (GWAS) and found evidence of contrasting locally adaptive clines between regions. Western Mediterranean ecotypes showed low δ^13^C/early flowering at low elevations to high δ^13^C/late flowering at high elevations, while Asian ecotypes showed the opposite pattern. We mapped different candidate genes for each region, and trait-associated SNPs often showed elevational clines likely maintained by selection. Antioxidants and pigmentation showed regional differentiation but rarely elevational clines. GWAS for antioxidants identified an ascorbate transporter *PHT4;4* (AT4G00370), which we show alters non-photochemical quenching kinetics under high light and may be involved in local adaptation to Moroccan mountains. The low-antioxidant *PHT4;4* GWAS allele was associated with lower *PHT4;4* expression and this haplotype was characterized by binding sites of a transcription factor family, DOF, involved in light response. Our results highlight how physiological and genomic elevational clines in different regions can be unique, underlining the complexity of local adaptation in widely distributed species.

## Introduction

Changes in environmental conditions with changing elevation are some of the most iconic natural gradients (Caldas, 1966). Correspondingly, changes in plant growth forms are dramatic over hundreds or thousands of meters of elevation (Caldas, 1966; Hedberg and Hedberg, 1979; Körner, 2021). Elevational gradients are among the key factor driving local adaptation, and hence diversity, within species that have broad elevational ranges (Clausen et al., 1940; Kooyers et al., 2015). At high elevations in particular, the combination of cold and high light during vegetative periods poses a major challenge. Additional environmental changes from sea level to alpine environments include wider fluctuations in daily temperature, decreased partial CO_2_ pressure (pCO_2_), altered (i.e., orographic) precipitation and clear sky radiation, and shorter growing seasons (Körner, 2003, 2007). While local adaptation to high elevation conditions has been detected in reciprocal transplants and common gardens (Villemereuil et al., 2018; Wos et al., 2022), its physiology and genetic basis is less clear.

Photosynthesis sits at the nexus of multiple physiological challenges for high elevation plants. For example, pCO_2_ declines approximately linearly with elevation, at 4000 m being only ∼65% of sea level pCO_2_ (Körner, 2021). Species from high elevations may exhibit higher fitness and better regulation of physiological processes under low atmospheric pCO_2_ (Ward and Strain, 1997; Zhu et al., 2010). However, pCO_2_ limitation is likely counteracted by elevational reductions in pO_2_ and photorespiration (Wang et al., 2017). Additionally, clear sky radiation increases with elevation, but cold temperatures slow the dark reactions of photosynthesis, leading to photo-oxidative damage if excess energy from photons is not properly handled (Wise, 1995). One mechanism to counteract photo-oxidative damage is non-photochemical quenching (NPQ) in which excess absorbed photons dissipate via the xanthophyll cycle (García-Plazaola et al., 2012; Muller et al., 2001). When light intensity rapidly increases (e.g., due to clearing cloud cover), rapid acclimation of NPQ (fast NPQ kinetics) may be adaptive (Rungrat et al., 2019). Another mechanism is the increased production of antioxidants (e.g., carotenoids and flavonoids), which can scavenge reactive-oxygen-species (Wise, 1995). Alpine environments can also experience extreme temperature fluctuations, often reaching below freezing temperatures every night, which not only exacerbates oxidative damage but can also physically damage cells (Suzuki and Mittler, 2006).

The model plant *Arabidopsis thaliana* (hereafter Arabidopsis) has a broad geographical, environmental, and elevational range, from below sea level to ∼4400 m, making it an ideal system for studying multi-regional elevational adaptation. Several studies have addressed elevational adaptation in Eurasia, though in single mountain systems. For example, in the Pyrenees low elevation/coastal sites experience early summer drought, potentially selecting for a drought-escape strategy (Montesinos-Navarro et al., 2011, 2012; Picó, 2012; Vidigal et al., 2016; Wolfe and Tonsor, 2014). In the Alps, flowering time and its gene expression may contribute to adaptation to seasonal freezing variation along elevation (Suter et al., 2014), but physiological clines are unclear (Günther et al., 2016; Lampei et al., 2019; Luo et al., 2015a). In the western Himalayas flowering time can be plastic in response to temperature, especially in ecotypes from high elevation (Singh and Roy, 2017). Little is known about elevational clines in African Arabidopsis, where populations are older than most Eurasian ones, have higher genetic diversity, and grow under the highest recorded elevations (up to ∼4400 m) in the unique Afroalpine ecosystem (Brennan et al., 2014; Brochmann et al., 2021; Durvasula et al., 2017).

One challenge in deriving general conclusions about adaptive responses to elevation for widespread species is that elevational changes in environment often differ from region to region, potentially leading to region-specific local adaptations. For example, while temperature consistently decreases with elevation, different regions may show different elevational clines in cloudiness, resulting in fluctuating light intensity that complicates elevational light patterns in different geographical regions (Körner, 2021). Even when gradients are consistent, local adaptation to isolated mountain ranges can occur via shared or distinct genetic and phenotypic mechanisms (Bohutínská et al., 2021). Furthermore, plant adaptation to extreme conditions often comprises complex multivariate responses where homeostatic regulation during photosynthesis is achieved in combination with changes in phenology (FernándezLJMarín et al., 2020; Kooyers et al., 2015; Smith and Knapp, 1990).

Here, we took a multi-regional view of Arabidopsis genomes, life history, and physiology across elevational gradients. We asked whether putatively adaptive physiological and genomic elevational clines were consistent among regions and hypothesized that ecotypes from higher elevations have robust performance in the face of high elevation stressors (i.e., better growth and resource use, lower oxidative stress) using a diverse set of naturally inbred lines (ecotypes). We integrate large-scale experiments and genome-wide association studies (GWAS) with detailed physiological panels, revealing the complexity of elevational adaptation across the range of Arabidopsis. For a GWAS candidate gene (*PHT4;4*), we explore the physiological effects of natural and mutant allelic variants in response to high light. With our regional approach in a widespread model plant, we show that different mountains exhibit different strategies of local adaptation along elevation.

## Results

### Population genetic structure and regional patterns in climate

To investigate natural phenotypic and genetic variation across elevational gradients in different geographical regions, we studied a total of 271 ecotypes, including 262 from native regions in Eurasia and Africa (Figure 1 A, Supplemental Dataset S1). We defined five geographical regions based on location and genetic cluster (Supplemental Figure S1): Asia (46 ecotypes), central Europe/Caucasus (59 ecotypes), northwestern Europe (26 ecotypes), western Mediterranean (112 ecotypes), and eastern Africa (19 ecotypes, including 9 sequenced). The last region encompasses the poorly studied genetically diverse Afroalpine Arabidopsis populations (Fulgione and Hancock, 2018).

**Figure 1.**
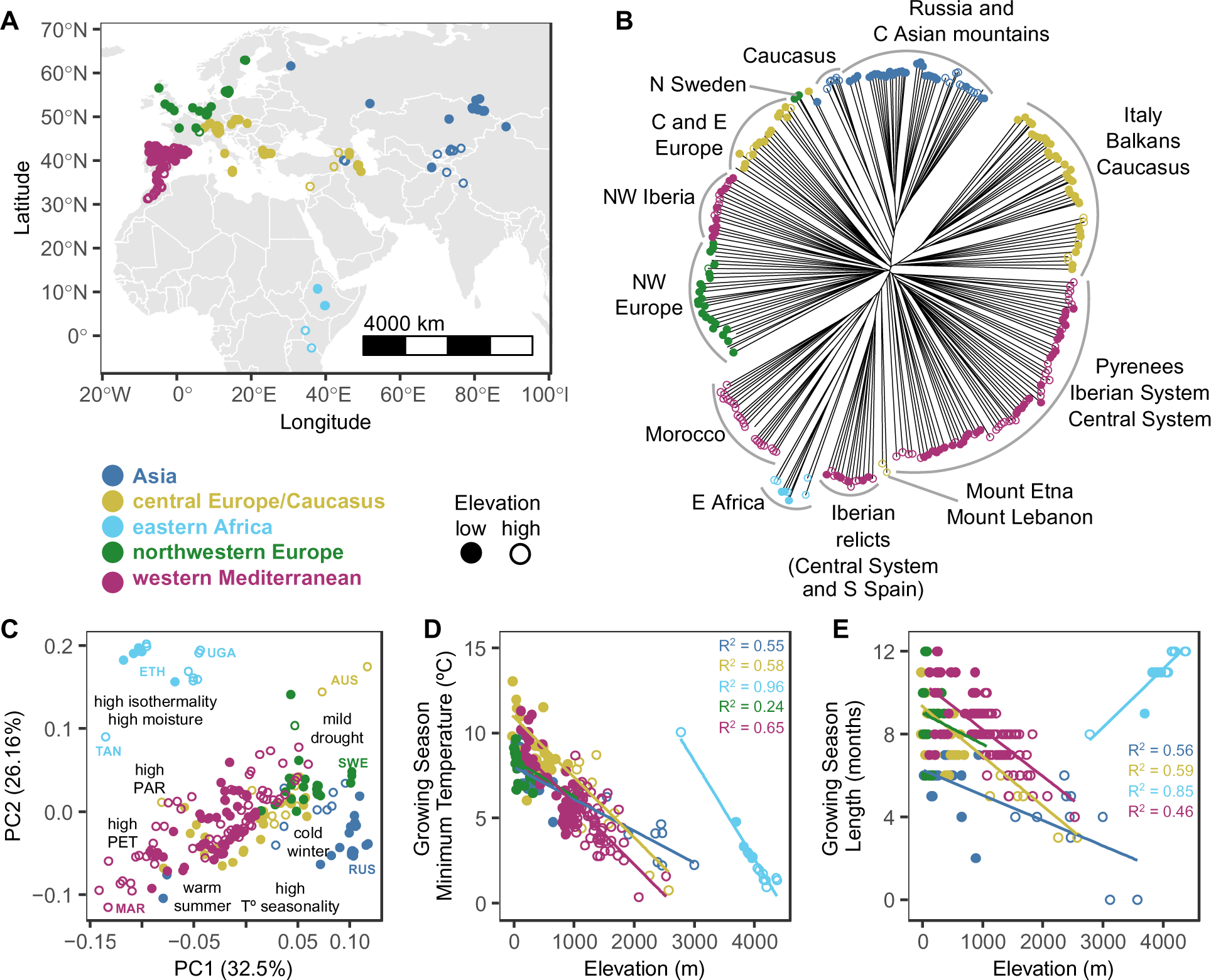
Provenance, genetic relationships, and climates of studied native ecotypes colored by region. Symbols represent elevation below (closed circles) and above (open circles) 1000 m for all regions, except for eastern Africa where the threshold is 4000 m. **A** Geographic location of ecotypes. **B** Neighbor-joining tree based on 261 ecotypes and 562,194 resequencing SNPs filtered for LD. Tips with no symbol are ecotypes outside of the native range. **C** PC1 and PC2 of climate space representing conditions in 257 ecotypes based on 85 climatic variables. Arrows and text summarize loadings more strongly correlated with PC1 and PC2, indicating the direction of the relationship (not proportional to eigenvectors). PAR=photosynthetically active radiation, PET=potential evapotranspiration, AUS=Austria, ETH=Ethiopia, MAR=Morocco, RUS=Russia, SWE=Sweden, TAN=Tanzania, UGA=Uganda. **D**, **E** Linear regressions of climatic variables on elevation of origin. Selected variables describe the growing season. Significant relationships (*p* < 0.01) are depicted by their R^2^.

We examined population genetic structure in our diversity panel using a published variant call format (VCF) file (Durvasula et al., 2017) that we combined with 16 ecotypes (eight phenotyped) from eastern Africa that we sequenced here (Supplemental Methods S1). We calculated pairwise genetic distances of 261 ecotypes (Supplemental Dataset S1, ‘sequenced’ column) from 562,194 whole genome resequencing single nucleotide polymorphisms (SNPs) filtered for minor allele frequency (MAF<0.05) and linkage-disequilibrium (LD). A neighbor-joining tree showed that population genetic structure largely corresponded to our geographical regions and not elevation. Ecotypes from low and high elevations on the same mountain range were more closely related than ecotypes of similar elevations on different mountains. In the Caucasus, however, ecotypes from two of our regions overlapped (Asia and central Europe/Caucasus), highlighting the genetic heterogeneity of this region (Figure 1 B).

A principal components analysis (PCA) of 85 climatic variables from the native range of 257 studied ecotypes with accurate provenance/climate data (Supplemental Dataset S2) showed that ecotypes from different regions occupied distinct climates and climatic extremes (Figure 1 C). Asia had prominent temperature seasonality, with short growing seasons and cold winters. Central Europe/Caucasus had milder temperature seasonality and warm growing seasons. Northwestern Europe was cold with mild dry seasons. In the western Mediterranean temperature variation was wide, with dry seasons at high elevation ranging from hot, dry, and bright (Atlas Mountains) to temperate and moist (Pyrenees). East Africa occupied the most distinct climate of long growing seasons with sharp day/night temperature changes. Elevational range and extent differed among regions, but environmental gradients were generally similar, with lower night temperatures and shorter growing seasons at higher elevations (Figure 1 D, E).

### Trait variation under low pCO2, high light and night-time freezing conditions

To characterize genetic variation in physiology, resource use, and performance under alpine-associated stressors (*i.e.*, trait differences among ecotypes in our diversity panel), we performed three separate experiments (12 h photoperiod) with subsets (111–254) of the 271 studied ecotypes (Supplemental Dataset S1). Subsets were different based on seed and space availability and germination success, with a total of 101 ecotypes overlapping between experiments. Our first experiment imposed low partial CO_2_ pressure (pCO_2_; with cool and bright conditions) on 253 ecotypes (Supplemental Dataset S3). Our next two experiments imposed contrasting light levels (250 vs 600 μmol m^−2^ s^−1^) on 111–114 ecotypes (performed simultaneously under cool conditions) and contrasting night-time temperatures (–4 vs 4°C) on 170–172 ecotypes (performed sequentially under bright conditions) (Supplemental Datasets S4–S5).

In our first experiment we measured δ^13^C and δ^15^N, two stable isotope ratios associated with carbon/water and nitrogen use, respectively, and integrated across the plant life span, root diameter, rosette diameter, and rosette compactness. We examined coordinated life history variation (*i.e.*, in life cycle and physiology) with a PCA of these traits along with published flowering time data (Alonso-Blanco et al., 2016) (N=206 ecotypes). Asian ecotypes tended to have smaller and more compact rosettes with smaller roots and higher δ^13^C and δ^15^N, in contrast to western Mediterranean ecotypes which showed greater phenotypic variation, including larger and less compact rosettes with larger roots lower δ^13^C and δ^15^N (Figure 2 A). For flowering time, we found high variation within regions, with generally later flowering in northwestern Europe (likely due to strict vernalization requirements), and earlier flowering in central Europe/Caucasus (likely due to weak flowering time cold response genes), while Asia and the western Mediterranean were not different to each other (Supplemental Figure S2 A). This high within-region diversity was also apparent at the level of some major known flowering time genes (Supplemental Figure S2 B).

**Figure 2.**
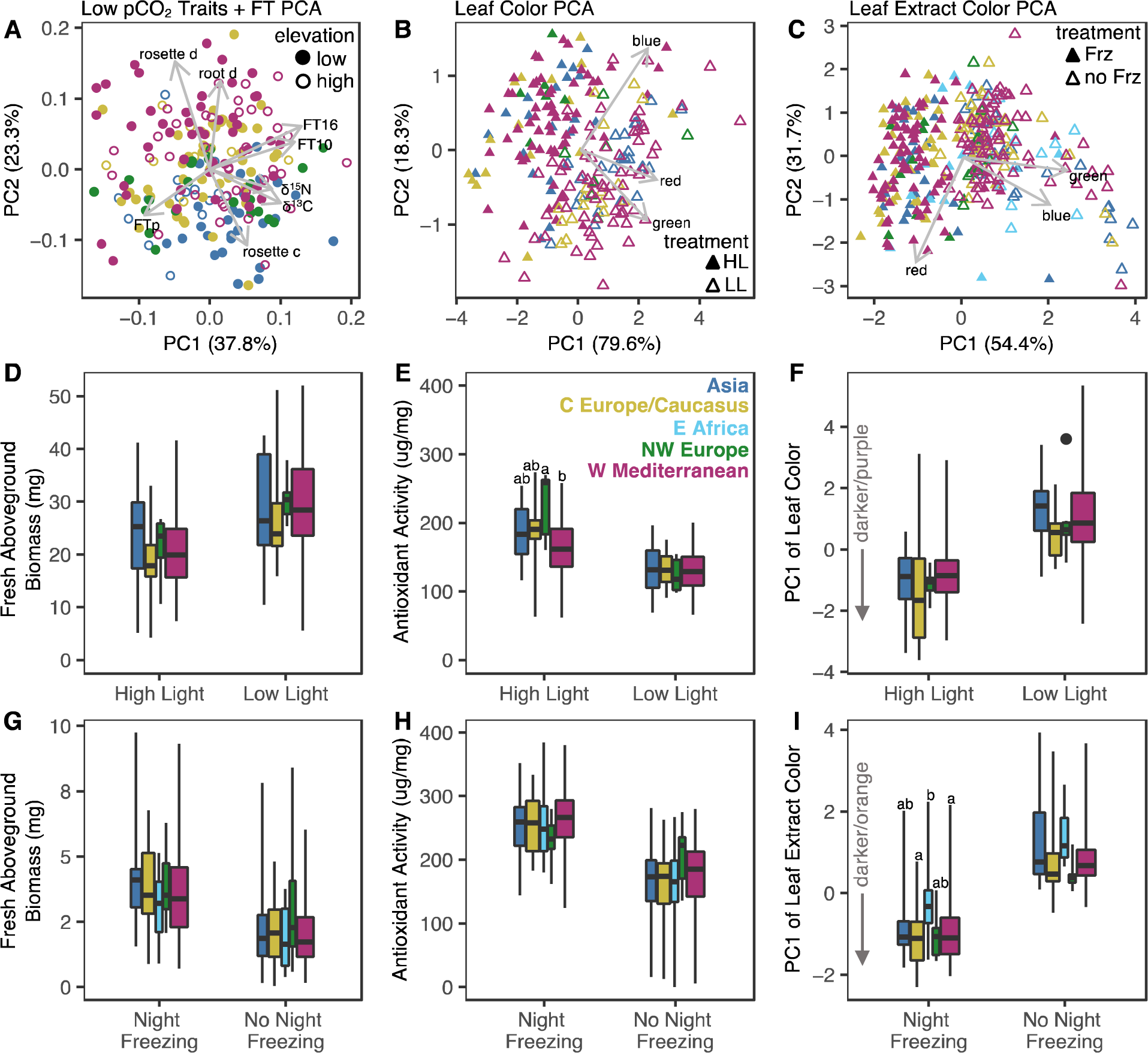
Ecotype responses to high elevation experimental conditions colored by region. **A** Eigenvector plot of the loadings of 8 variables (d: diameter, c: compactness) onto PC1 and PC2 of phenotypes under low pCO_2_ along with flowering time (FT at 10°C: FT10 and at 16°C: FT16) and flowering time plasticity (FTp: FT10–FT16) in 206 ecotypes (5–6 replicates). **B–C** PC1 and PC2 of RGB colors in experiment 2 (from rosette photos of 110 ecotypes-5 replicates under high light: HL vs. low light: LL) and experiment 3 (from leaf-extract photos of 163 ecotypes-6 pooled replicates under night-time freezing: Frz vs. no night-time freezing: no Frz), respectively. **D–I** Significant phenotypic plasticity under contrasting light (center: Asia N=23, C Europe/Caucasus N=17, NW Europe N=7, W Mediterranean N=63) and night-time temperatures (bottom: Asia N= 27, C Europe/Caucasus N=38, NW Europe N= 10, E Africa N=13, W Mediterranean N=75). Boxplots are proportional to sample size and depict the median and interquartile range, whiskers cover the data extent. **D, G** Fresh aboveground biomass (mg). **E, H** Antioxidant activity (ug per leaf mg). **F, I** PC1 of RGB color from leaves and leaf extracts, respectively.

In our next experiments, we found significant trait changes between treatments and trait differences among regions (ANOVA, *p*<8E–10; Figure 2) in fresh aboveground biomass, total antioxidant activity, and the multidimensional color variation of leaves (high light experiment) and their extracts (night-time freezing experiment) based on a PCA of RGB color values (Figure 2 B, C, Supplemental Results S1). Color may indicate phenotypes connected to elevational adaptation because pigments can be adaptive as they act as antioxidants under high light and/or cold (Havaux and Kloppstech, 2001).

Under cool temperatures and high light, plants were smaller, had higher antioxidant activity, and dark purple (vs. green) leaves (lower values of PC1_LeafColor_/higher values of PC2_LeafColor_) than when grown under low light (Figure 2 B, D–F). Survival rates were slightly but significantly lower under high than low light (93 vs 96%, *p*=0.02), and positively correlated with rosette biomass (Pearson correlation coefficient (cor)=0.23, *p*=0.01) under high light. Furthermore, ANOVA showed significant differences in antioxidant activity at high light among regions (*p*=0.004), with western Mediterranean ecotypes having the lowest values (Figure 2 E), suggesting lower sensitivity to high light.

Under night-time freezing, plants were larger, had higher antioxidant activity, and more orange (as opposed to yellow) leaf extracts (lower values of PC1_ExtractColor_) than when grown with night-time temperatures just above freezing (Figure 2 C, G–I). Survival was significantly lower under night-time freezing than under no freezing (76 vs 100%, *p*<0.0001), and in the western Mediterranean survival was negatively correlated with higher antioxidants under night freezing (*i.e.*, antioxidant under night freezing - antioxidant under no freezing; cor= –0.38, *p*=0.001), suggesting maladaptive plasticity. Additionally, ANOVA showed significant differences in PC1_ExtractColor_ under night-time freezing between regions (i.e., orange in extracts, *p*=0.007), with eastern Africa ecotypes (all from >2775 m) having yellower extracts (Figure 2 I). In eastern Africa more orange extracts (lower values of PC1_ExtractColor_) were associated with higher survival rates (cor= –0.92, *p*=0.001, df=6), suggesting selection associated with pigments.

Taken together, ecotypes showed great variation in flowering time and in response to high elevation experimental conditions. Ecotypes from different geographical regions showed distinct morphology under low pCO_2_ (bright and cool conditions) and strong treatment effects on all phenotypes measured under high vs. low light, and under night-time freezing vs. night-time above freezing. We detected some differences in phenotypes among regions, suggesting decreased sensitivity in antioxidants to high light in western Mediterranean ecotypes (Figure 2 E), and decreased content in leaf extract greenness under night-time freezing in eastern African ecotypes (Figure 2 I).

### Genetic variation in NPQ kinetics under cold and drought

Non-photochemical quenching (NPQ) kinetics in response to fluctuating light (e.g., due to cloud changes) can be an important mechanism for limiting photooxidative damage at high elevations (Kumar et al., 2021). To capture diverse strategies, we selected a global subset of 11 ecotypes representing the lowest (22–254 m) and highest (1576–4078 m) elevational range of Arabidopsis (Supplemental Dataset S1). We performed detailed measurements of NPQ kinetics under fluctuating light-dark cycles in response to two potential high elevation stressors: cold (22 vs. 4°C under well-watered conditions for 17 h) and drought (daily-watering vs. 5-day-drought under 22°C). We fitted exponential equations to individual NPQ time series to obtain parameters that describe NPQ induction and relaxation (Sahay et al., 2023, Supplemental Figure S3).

We found that cold and drought significantly affected NPQ induction and relaxation, but this effect significantly varied between ecotypes (ANOVA, Supplemental Table S1, Supplemental Figures S4–S5). For example, CYR from low and seasonally dry France showed a slow NPQ response to cold but a fast response to drought via rapid NPQ induction (Figure 3 A–B). Conversely, Dja-1 from high and cold Kyrgyzstan showed no changes in NPQ kinetics in response to drought but a strong response to cold via sustained high NPQ after fluctuating light treatments (2^nd^ and 3^rd^ cycles; Figure 3 C–D), which could be important to avoid high light stress under cold. Sij-1 from high and cold Uzbekistan showed a similar pattern (Figure 3 G–H). In IP-Trs-0 from low and mesic Iberia, cold and drought had similarly minor effects on NPQ kinetics (Figure 3 E–F), suggesting reduced sensitivity to these abiotic stressors. Interestingly, under control conditions, IP-Trs-0 and JL011-2-1, an ecotype with Afroalpine origin where nights are subject to below freezing temperatures all-year-round, had the lowest fresh weight and the highest dry matter content (>60%), indicating low water content (Supplemental Figure S6), which could aid in drought-avoidance or in freezing-tolerance. Additionally, under control conditions, JL011-2-1 had the lowest chlorophyll and total carotenoid content, but the highest chlorophyll (*a* + *b*) to carotenoids ratio (Supplemental Table S2).

**Figure 3.**
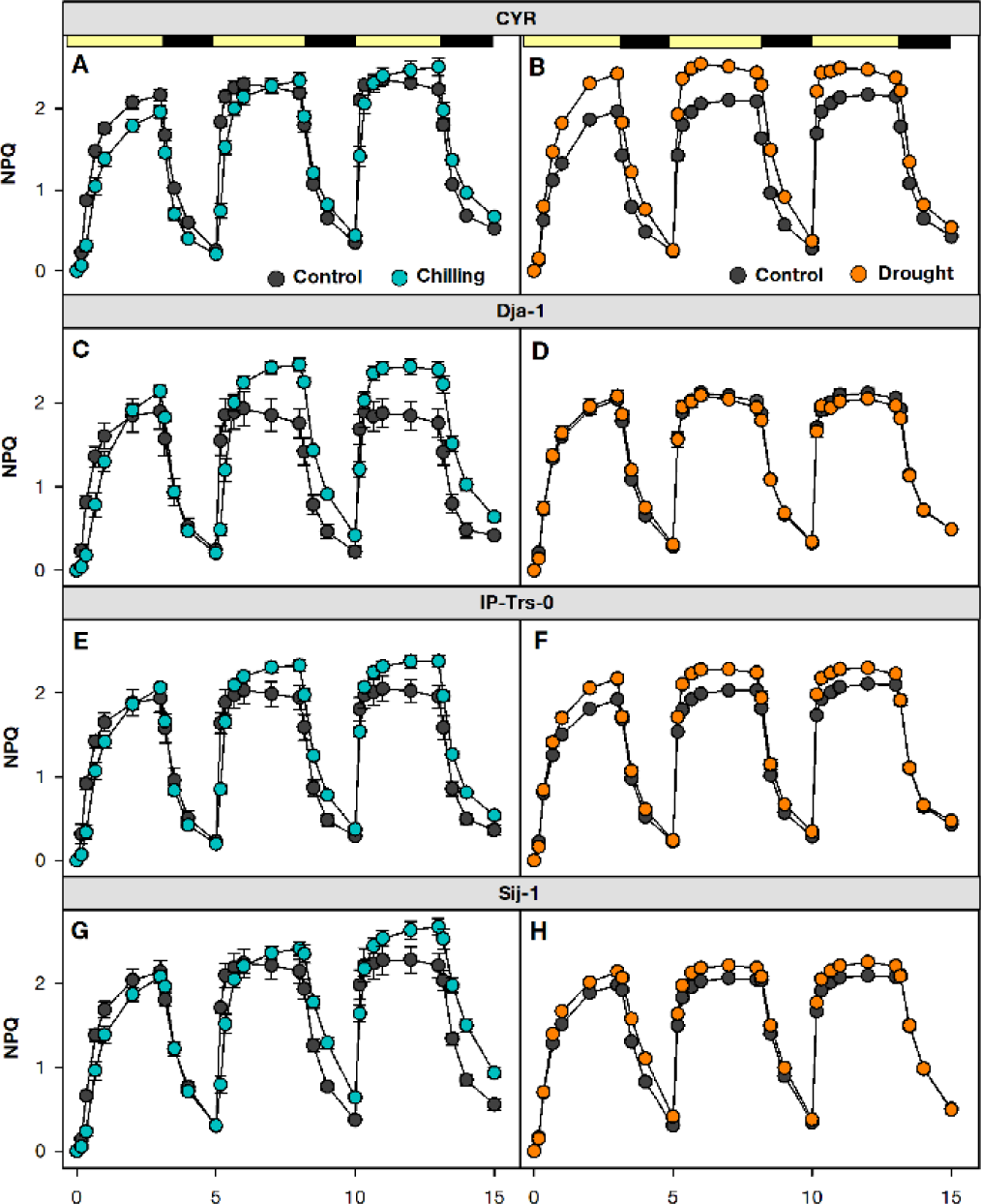
Four representative examples of non-photochemical quenching curves during three sequential light-dark cycles under cold (left panels) and drought (right panels) with corresponding control. Chosen ecotypes represent the lowest and highest elevations of the Arabidopsis range: **A**–**B** CYR (52 m). **C–D**. Dja-1 (2995 m). **E–F** IP-Trs-0 (254 m). **G–H** Sij-1 (2459 m). Bars are means ± SE (5–10 replicates).

Taken together, while there was no clear pattern relative to elevation of origin across this global sample of ecotypes, responses to drought or cold in NPQ kinetics greatly varied among ecotypes, with responses potentially corresponding to the environmental pressures at their sites of origin.

### Regional elevational clines in phenotype and the genomic basis of phenotypic variation

To understand how different elevational gradients shape phenotypic (Figure 4) and genomic variation, we tested for region-specific trait- and SNP-elevation clines with linear mixed models that controlled for genome-wide similarity among ecotypes (kinship), which when significant suggest that selection is driving clines (Josephs et al., 2019). GWAS that controlled for kinship (Supplemental Datasets S6–S9) were performed on our global dataset of 19 phenotypes (N=110– 253 ecotypes, 2.7–2.8M SNPs), on regional datasets for six phenotypes that significantly varied with elevation (N=44–112, 1.9–2.8M SNPs), and on regional datasets for flowering time at 10 and 16°C and flowering time plasticity (*i.e.*, flowering time at 10°C minus flowering time at 16 °C; N=67–398, 2–2.6M SNPs). We report candidate protein coding genes whose annotated function relates to phenotype function (Table 1) and highlight regional QTL for which top SNP significantly changed with elevation in mixed models that controlled for kinship.

**Figure 4.**
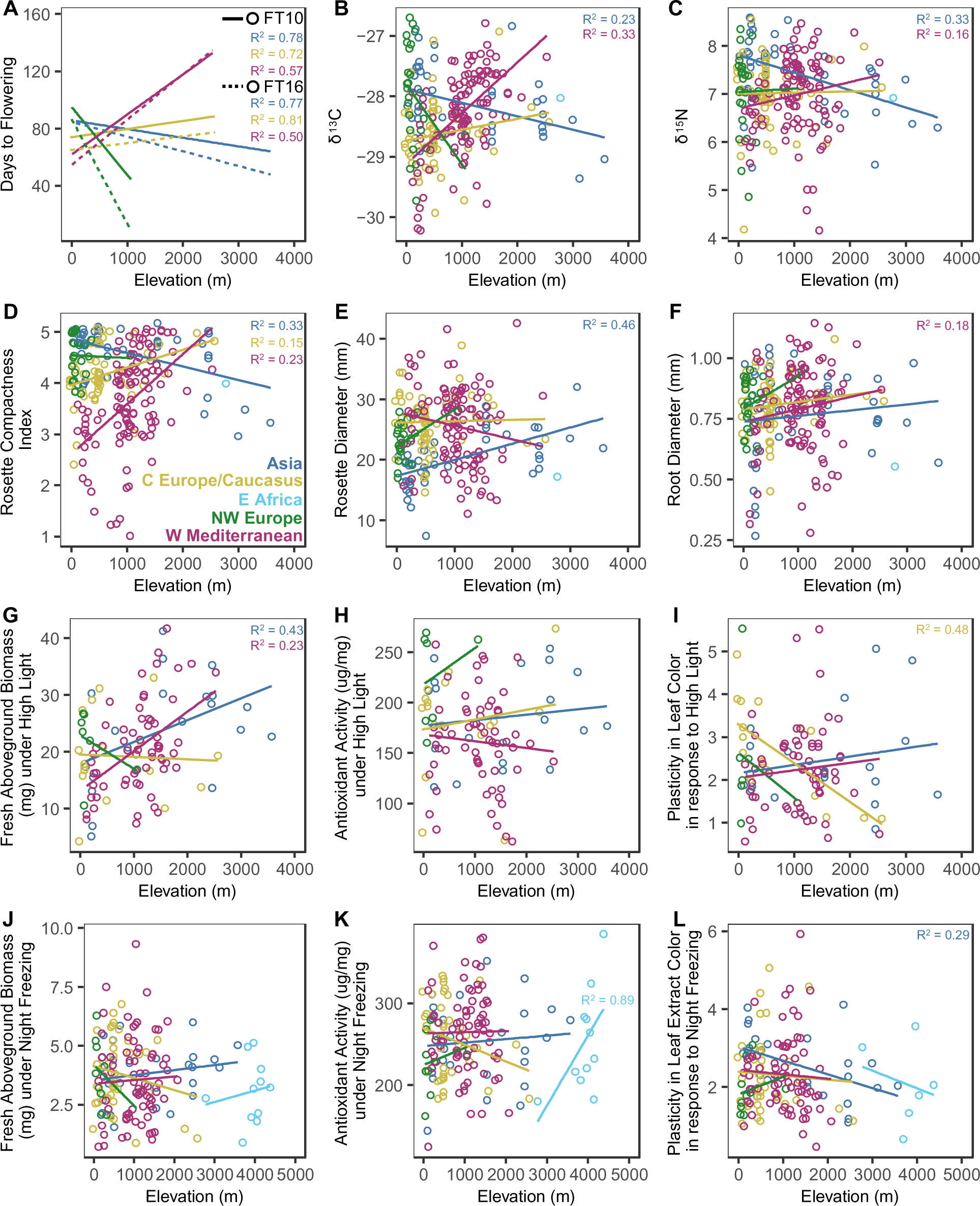
Linear regressions on elevation of origin of flowering time (FT) and phenotypes under low pCO_2_, high light, and night-time freezing. Ecotypes and fitted lines are colored by region: Asia N=20–60, central Europe/Caucasus N=17–264, eastern Africa N=5–11, northwestern Europe N=7–331, and western Mediterranean N=64–174. Significant relationships from linear mixed effects kinship models (p<0.05) are depicted by their likelihood-ratio based pseudo-R^2^. **A** Days to flower at 10°C (FT10) and 16°C (FT16) obtained from the 1001 Genomes Consortium. **B**–**F** Phenotypes measured in a low pCO_2_ growth chamber. **G–I** Selected phenotypes measured in a high light growth chamber. J–L. Selected phenotypes measured in a night-time freezing growth chamber.

**Table 1.** Candidate genes mapped with GWAS which associated SNP follows a putatively adaptive elevational cline. CHR: chromosome, Coord: genomic coordinates in bp, Rosette D: rosette diameter, Rosette Comp: rosette compactness, W Med: western Mediterranean.

| Locus | Gene | Function | CHR | Coord (bp) | Strand | SNP (bp) | Phenotype | Region |
| --- | --- | --- | --- | --- | --- | --- | --- | --- |
| AT1G77080 | *FLOWERING LOCUS M* | Flowering response to cold | 1 | 28955522–28960111 | Forward | 28957836 | $\delta^{13}\text{C}$ | Asia |
| AT1G65490 | *SECRETED PEPTIDE STMP5* | Plant growth and pathogen defense | 1 | 24354646–24355490 | Forward | 24356159 | Rosette D | Asia |
| AT5G41890 | GDSL-motif | Biotic and abiotic stress responses | 5 | 16764211–16766702 | Reverse | 16766371 | Rosette Comp | Asia |
| AT5G45830 | *DELAY OF GERMINATION1* | Seed dormancy response to cold | 5 | 18589438–18591506 | Reverse | 18590591 | FT10 & FT16 | W Med |
| AT5G23220 | *NICOTINAMID-ASE 3* | Absciscic acid signaling | 5 | 7819608–7820760 | Reverse | 7818730 | $\delta^{13}\text{C}$ | W Med |
| AT5G11850 | MAP3 kinase | Absciscic acid signaling | 5 | 3816080–3821170 | Reverse | 3821491 | $\delta^{13}\text{C}$ | W Med |
| AT4G00370 | Phosphate transporter (*PHT4;4*) | Ascorbate transporter | 4 | 162897–166357 | Reverse | 160735 | HL antioxidants | Global |
| AT5G49730 | *FERRIC REDUCTION OXIDASE 6* | Iron transporter | 5 | 20201037–20204595 | Reverse | 20206030 | HL antioxidants | Global |
| AT5G49740 | *FERRIC REDUCTION OXIDASE 7* | Iron transporter | 5 | 20205302–20208777 | Reverse | 20206030 | HL antioxidants | Global |

#### Evidence of adaptive elevational clines in Asia

Using mixed models that accounted for kinship among ecotypes we found evidence that several traits showed elevational clines maintained by selection. Flowering time (at 10 and 16°C), δ^13^C, δ^15^N, rosette compactness, and plasticity in leaf extract color in response to night-time freezing (*i.e.*, the Euclidean distance in PC 1-2 space between contrasting conditions for a given ecotype, Figure 2 C), decreased with elevation (all *p*<0.03). Conversely, flowering time plasticity, rosette diameter, stomatal density, biomass under high and low light, and survival rates under high light and night-time freezing, increased with elevation (all *p*<0.025; Figure 4 A–G). We found corroborating evidence in our growth chamber experiments. Under high light stress, flowering time (at 10 and 16°C) was negatively correlated with both biomass and survival (cor ranged from –0.45 to –0.59, all *p*<0.04), while δ^13^C was negatively correlated with survival (cor= –0.46, *p*=0.03). Thus, high elevation ecotypes from this region seemingly showed a life history strategy of rapid phenology but inefficient water and nitrogen use, where accelerated reproduction and low water use efficiency (WUE) under high light stress may enhance biomass and survival.

GWAS mapped quantitative trait loci (QTL) for δ^13^C, rosette compactness, and rosette diameter, that showed elevational variation potentially maintained by selection (*i.e.*, after controlling for kinship, p<0.05), although for δ^13^C and rosette diameter no SNP passed FDR<0.05 in the GWAS (Figure 5 A, D, G). For δ^13^C, the 25^th^ top QTL was *FLOWERING LOCUS M* (*FLM*; AT1G77080), a negative regulator of flowering in response to cold, where loss-of-function removes the response to cool (≤10°C) temperatures and results in earlier flowering (Jin et al., 2022). For rosette compactness, the top QTL was a GDSL-like protein (AT5G41890) which is poorly studied and belongs to a family of lipolytic enzymes involved in morphogenesis and development (Lai et al., 2017). For rosette diameter, the 3^rd^ top QTL was downstream of a secreted peptide (*STMP5*; AT1G65490) that functions in plant growth and potentially disease defense (Yu et al., 2019).

**Figure 5.**
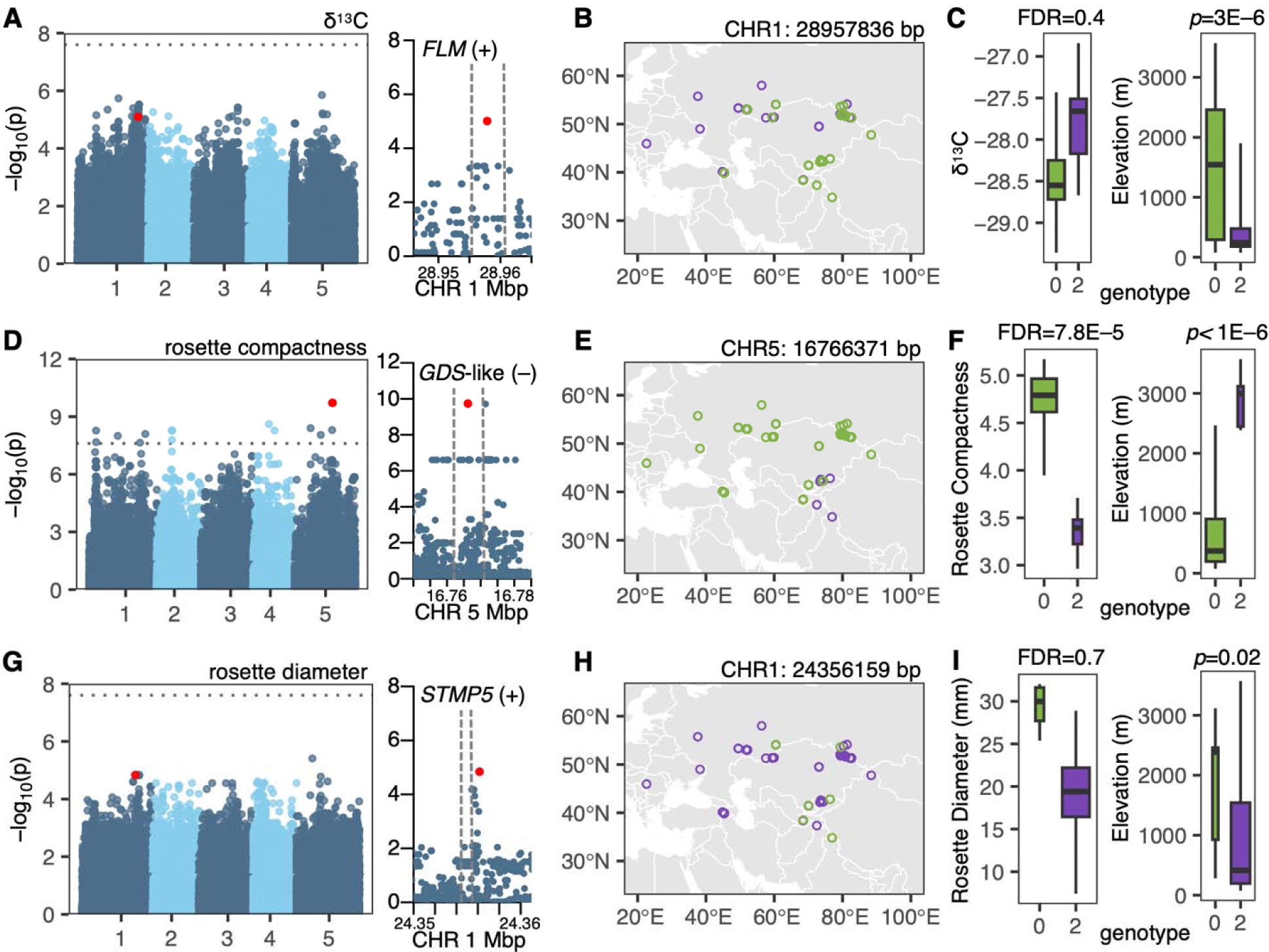
Genome-wide association studies (GWAS), geographic distribution, and variation of GWAS SNP alleles for selected traits and associated SNPs that showed elevational variation in Asia: δ^13^C (top panels), rosette compactness (middle panels), and rosette diameter (bottom panels). **A**, **D**, **G** Manhattan plots over the five chromosomes of the Arabidopsis nuclear genome with hit SNPs marked in red and dotted lines corresponding to a genome-wide Bonferroni threshold. Right panels are zoomed-in on candidate gene QTL (surrounded by dashed lines), showing their orientation (+: forward, –: reverse) and associated SNP marked in red. SNPs were filtered for MAF=0.05, N=45 ecotypes and 1,984,688 SNPs for δ^13^C and N=46 ecotypes and 2,002,562 SNPs for rosette compactness and diameter. **B**, **E**, **H** Map of GWAS SNP alleles in Asia. **C**, **F**, **I** Variation of GWAS SNP alleles (0: alternate allele, 2: reference allele) relative to traits of interest (left, with GWAS FDR), and elevation (right, with *p*-value from a kinship mixed model). Boxplot width is proportional to allele frequency and depict the median and interquartile range, whiskers extend to the data extremes.

Allelic variants in the aforementioned candidate genes involved in high δ^13^C, and compact/small rosettes, were restricted to low elevations (linear-mixed kinship model *p*=0.000003 for chr. 1: 28957836 bp, *p*<0.000001 for chr. 5: 16766371 bp, *p*=0.02 for chr. 1: 24356159 bp, respectively), suggesting that changing selection with elevation on *FLM*, the GDSL-like enzyme, and *STMP5* expression contributes to elevational clines in δ^13^C via flowering time variation in response to cold and in rosette morphology via growth and development. Although these SNPs were not found among the top 100 in our flowering time GWAS, the high δ^13^C/rosette compactness variants were significantly associated with delayed flowering at 10°C (Wilcox-tests *p*=0.002 for chr. 1: 28957836 bp and *p*=0.003 for chr. 5: 16766371 bp).

#### Evidence of adaptive elevational clines in the western Mediterranean

We found evidence for clines maintained by selection in flowering time (at 10 and 16°C), δ^13^C, δ^15^N, root diameter, rosette compactness, and biomass under high and low light, which all increased with elevation (all *p*≤0.03; Figure 4 A–G). Additionally, stomatal density was positively correlated with δ^13^C (cor=0.39, *p*=0.02) and flowering time (at 10 and 16°C: cor=0.49 and 0.45, *p*=0.007 and 0.02, respectively), suggesting that stomatal patterning varies with life history and water use strategies. Thus, high elevation ecotypes from this region apparently displayed a strategy of delayed phenology and efficient water and nitrogen use, directed toward root and aboveground biomass accumulation.

We found elevational variation in GWAS QTL for flowering time and δ^13^C (all FDR<0.05; Figure 6). For flowering time, the top QTL (at 16°C) and the 2^nd^ top QTL (at 10°C) was *DELAY OF GERMINATION 1* (*DOG1*; AT5G45830), an extensively studied gene involved in regulating seed dormancy and flowering time (Alonso-Blanco et al., 2016; Graeber et al., 2014; Huo et al., 2016; Martínez-Berdeja et al., 2020). For δ^13^C, the top and the 7^th^ top QTL were adjacent (downstream and upstream, respectively) of two genes involved in regulation of abscisic acid (ABA) in response to osmotic stress. The first is *NICOTINAMIDASE 3* (*NIC3*; AT5G23220), which encodes an enzyme that prevents the accumulation of intracellular nicotinamide by converting it into nicotinic acid in the NAD+ pathway, and which is expressed via Repressor of Silencing 1 (ROS1) DNA demethylation in response to ABA (Kim et al., 2019). The other is *MAP3 kinase* (*MAP3k*; AT5G11850), which is involved in the phosphorylation of SnRK2 kinases that in turn trigger osmotic-stress and ABA responses (Takahashi et al., 2020).

**Figure 6.**
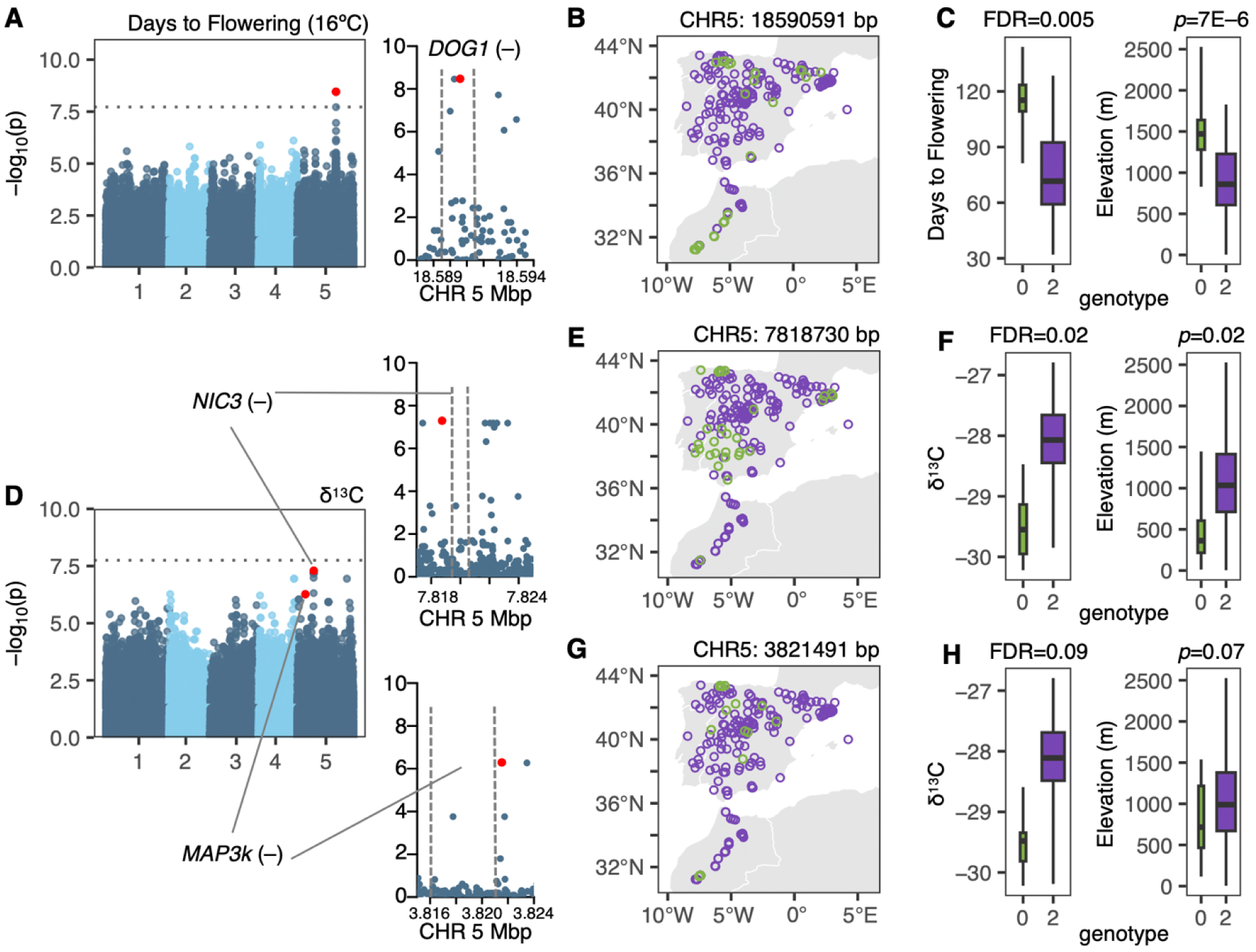
Genome-wide association studies (GWAS), geographic distribution, and variation of GWAS SNP alleles for selected traits and associated SNPs that showed elevational variation in the western Mediterranean: flowering time (top panels) and δ^13^C (middle and bottom panels). **A**, **D** Manhattan plots over the five chromosomes of the Arabidopsis nuclear genome with hit SNPs marked in red and dotted lines corresponding to a genome-wide Bonferroni threshold. Right panels are zoomed-in on candidate gene QTL (surrounded by dashed lines), showing their orientation (+: forward, –: reverse) and associated SNP marked in red. SNPs were filtered for MAF=0.05, N=178 ecotypes and 2,663,439 SNPs for flowering time (at 16°C) and N=110 ecotypes and 2,831,054 SNPs for δ^13^C. **B**, **E**, **G** Map of GWAS SNP alleles in the western Mediterranean. **C**, **F**, **H** Variation of GWAS SNP alleles (0: alternate allele, 2: reference allele) relative to traits of interest (left, with GWAS FDR), and elevation (right, with *p*-value from a kinship mixed model). Boxplot width is proportional to allele frequency and depict the median and interquartile range, whiskers extend to the data extremes.

SNP alleles involved in late flowering and high δ^13^C were restricted to high elevation (linear-mixed kinship model *p*=0.000007 for chr5: 18590591 bp and *p*=0.02 for chr5: 7818730 bp, although *p*=0.07 for chr5: 3821491 bp, respectively), suggesting changing selection with elevation on *DOG1*, *NIC3* and *MAP3k* contributes to elevational clines in flowering time via seed dormancy variation and in δ^13^C via the ABA pathway and stomatal dynamics. In Iberia, allelic variants associated with low δ^13^C/rapid flowering occurred in coastal and lowland areas with late-spring heat and drought (Figure 6), where ecotypes appear to display a drought-escape strategy (Montesinos-Navarro et al., 2011; Wolfe and Tonsor, 2014).

The *DOG1* SNP (chr5: 18590591 bp) was previously identified by Martínez-Berdeja et al. (2020) because of its association with seed dormancy in response to cold. The late flowering allele tags some (15 of 52 in Iberia and 15 of 68 in Morocco) of the ecotypes carrying the widespread ancestral haplotype of the *DOG1* self-binding domain (ECCY), but this SNP did not segregate in ecotypes with other haplotypes, nor in ECCY ecotypes from Asia (which only carry the other GWAS SNP allele). Delayed flowering was more frequent in ecotypes carrying the ECCY haplotype in all regions except in Asia (Kruskal-Wallis *p*=0.8 in Asia and *p*<0.0001 for all other regions; Supplemental Figure S7). Ecotypes carrying ECCY behave as winter annuals: seeds germinate soon after dispersal in the late summer–early fall (if not, extended cold induces secondary dormancy), plants overwinter as small rosettes, and flower over the spring (i.e., delayed flowering after vernalization) (Martínez-Berdeja et al., 2020). Our finding indicates that in Asia, delayed flowering is not exclusive to ecotypes carrying the ECCY overwintering haplotype, but also occurs in ecotypes carrying the D-SY haplotype where seeds only germinate after cold exposure (Martínez-Berdeja et al., 2020).

Martínez-Berdeja (2020) noted global climatic associations between *DOG1* haplotypes, including elevation, but did not characterize the direction of elevational clines. Here we found that *DOG1* haplotype frequency significantly changes with elevation in all regions (Kruskal-Wallis *p*<0.01), except in central Europe/Caucasus (*p*=0.3; Supplemental Figure S7). Relative to other haplotypes, ECCY was associated with higher elevations in the western Mediterranean, but with lower elevations in Asia and northwestern Europe. In Asia, the more frequent D-SY *DOG1* haplotype (Supplemental Figure S7) or mutations at other genes could also confer delayed flowering, which is the low elevation strategy in this region (opposite the western Mediterranean). Thus, the role of *DOG1* in local adaptation along elevation likely differs from region to region due to alternate life-history clines among regions.

#### Latitudinal but not elevational clines detected in northwestern Europe

We found no significant trait-elevational clines (all FDR>0.05). However, the elevational gradient in this group was narrow, occupying the lowest part of the range: 0–500 m, with only one ecotype at 1060 m. Similar to the latter two regions, flowering time (at 10 and 16°C) was positively correlated with δ^13^C and δ^15^N (cor=0.52–0.68, *p*=0.01–0.0006), while negatively correlated with rosette diameter (cor= –0.50 and –0.45, *p*= 0.02 and 0.04, respectively). We examined changes in phenotype along latitude, given that winters become longer and colder at higher latitudes, which may in turn relate to cold stress response. We found that delayed flowering and high δ^13^C were associated with higher latitudes (*p*<0.0001) in mixed models controlling for kinship among ecotypes, suggesting that overwintering at higher latitudes is adaptive via delayed flowering and high-water use efficiency. We did not perform GWAS on measured phenotypes for this region, given their small sample size (up to 26), but report flowering time QTL (see Supplemental Results S2).

#### Potentially adaptive elevational clines in central Europe/Caucasus but no clear QTL

Mixed models that accounted for kinship among ecotypes showed that flowering time (at 10 and 16°C) and rosette compactness increased with elevation as in the western Mediterranean (all *p*<0.05; Figure 4 A, D). Stable isotopes did not vary with elevation in this region, but as in other regions both δ^13^C and δ^15^N were positively correlated to flowering time at 10 and 16°C (cor=0.31–0.41, *p*=0.01–0.001). Although flowering time was not correlated to leaf color traits under stress, plasticity in leaf color in response to high light was lower (i.e., lower photosensitivity) at higher elevations (linear-mixed model, *p*=0.0001, Figure 4 I). We did not detect QTL mapped with GWAS that seemed functionally relevant for rosette compactness, and report one flowering time QTL that was, however, not associated with elevation (see Supplemental Results S2).

#### Elevational clines in eastern Africa

Afroalpine ecotypes were only included in the night-time freezing experiment and represented the highest part of the Arabidopsis range (mostly 3691– 4374 m, with one ecotype at 2775 m). Although based on a limited sample size of seven ecotypes, unlike other regions, here a mixed model that accounted for kinship found that antioxidant activity under night-time freezing increases with elevation (*p*=0.005; Figure 4 K).

#### Genotypes and phenotypes of montane Moroccan ecotypes under high light

In our global dataset, we mapped two QTL associated with antioxidant activity under high light stress (Figure 7 A). The top tagged *Ferric Reduction Oxidase 7* (*FRO7*; AT5G49740) and was upstream (in the putative promoter region) of *FRO6* (AT5G49730), both involved in iron chloroplast uptake and expressed in aerial green tissue (Jain et al., 2014; Jeong et al., 2008, 2009). *FRO6* expression is light-dependent, with several light-responsive elements in the promoter region (Feng et al., 2006). The second top QTL was downstream of a phosphate transporter (*PHT4;4*; AT4G00370) that carries ascorbate into the chloroplast (Miyaji et al., 2015). *PHT4;4* expression is higher in the chloroplast envelope membrane and increases under high light stress, with knockouts having lower levels of reduced ascorbate (an antioxidant) in the leaves, and lower content of ascorbate-dependent xanthophylls and ß-carotene after high light exposure (Miyaji et al., 2015).

**Figure 7.**
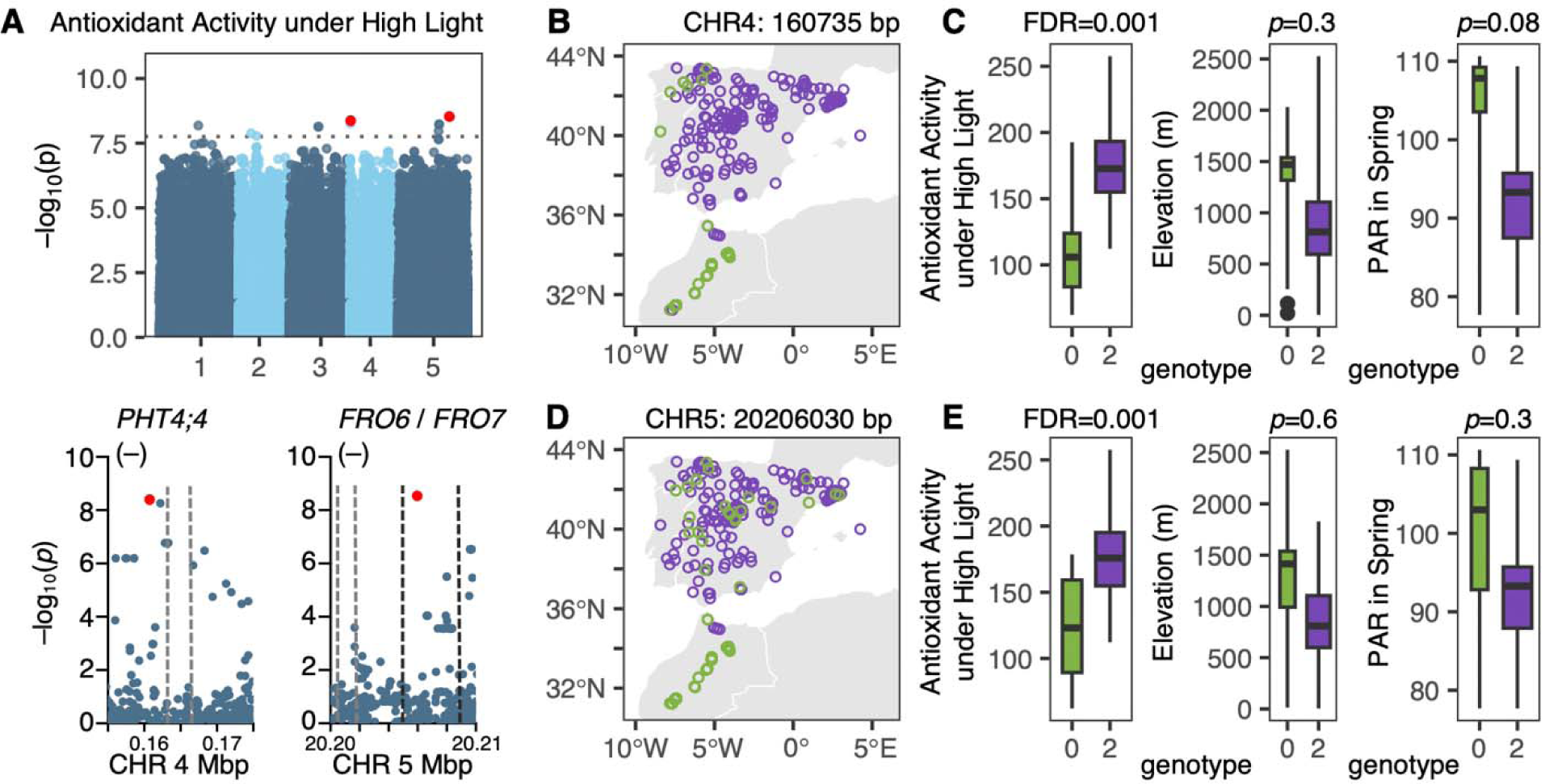
Genome-wide association study (GWAS), geographic distribution, and variation of GWAS SNP alleles associated with antioxidant activity under high light. **A** Manhattan plots over the five chromosomes of the Arabidopsis nuclear genome with hit SNPs marked in red and dotted lines corresponding to a genome-wide Bonferroni threshold. Bottom panels are zoomed-in on candidate gene QTL (surrounded by dashed lines), showing their orientation (+: forward, –: reverse) and associated SNP marked in red. SNPs were filtered for MAF=0.05, N=110 ecotypes and 2,865,792 SNPs. **B**, **D** Map of GWAS SNP alleles in the western Mediterranean. **C**, **E** Variation of GWAS SNP alleles (0: alternate allele, 2: reference allele) in the western Mediterranean relative to antioxidant activity under high light (left, with GWAS FDR), elevation (center, with *p*-value from a kinship mixed model), and spring photosynthetically active radiation (PAR) (left, with *p*-value from a kinship mixed model). Boxplot width is proportional to allele frequency and depict the median and interquartile range, whiskers extend to the data extremes.

To account for potential under-fitting of the genomic background effect (kinship random effects) for the genetically distinct Moroccan ecotypes (Atlas Mountains in Figure 1 B) that also had low antioxidant activity (Figure 2 E, 4 H), we performed another GWAS on this trait incorporating a covariate that indicated if the accession was from Morocco. P-values on this GWAS were higher (FDR>0.9) and previously mapped QTL were no longer mapped. The original GWAS results were thus driven by the low antioxidant activity of most Moroccan ecotypes.

Variants in top SNPs (chr5: 20206030 bp for *FRO6/FRO7* and chr4: 160735 bp for *PHT4;4*) associated with low antioxidant activity were globally uncommon (allele frequency AF=0.09 and 0.08, respectively), but dominant (AF=0.81 and 0.84) in montane (>1000 m) Moroccan ecotypes, and infrequent (AF=0.13 for chr5: 20206030 bp) to rare (AF=0.03 for chr4: 160735 bp) in Iberian ecotypes. Furthermore, these SNPs were strongly linked in Morocco (cor=0.36, *p*=0.004), but not elsewhere (cor= –0.01, *p*=0.8). In the western Mediterranean, the respective low-antioxidant GWAS SNP alleles were more frequent in ecotypes from sites with higher photosynthetically active radiation during the growing season (Kruskal-Wallis *p*=0.0001 for chr4: 160735 bp and Kruskal-Wallis *p*=0.003 for chr5: 20206030 bp), but only for chr4: 160735 bp the low-antioxidant allele was dominant at higher elevations (Kruskal-Wallis *p*= 0.004), but these patterns were not significant in mixed-linear kinship models (Figure 7 C, E).

To further assess evidence of selection on these loci, we calculated F_ST_ values for all SNPs with MAF>0.05 in chromosome 4 (where *PHT4;4* is located) and 5 (where *FRO6/FRO7* are located), comparing Moroccan ecotypes to the global panel. We found F_ST_=0.19 for chr4: 160735 bp and was in the 0.005 tail for the SNPs on the chromosome, and Chr 5: 20206030 had a similar pattern: F_ST_=0.14 and was in the 0.01 tail for the SNPs on the chromosome, suggesting these variants might be locally adaptive in Morocco.

Using published transcriptome data on 727 diverse ecotypes under typical low light laboratory conditions (Klepikova et al., 2016), we found that *PHT4;4* expression was weakly associated with allelic variation at chr4: 160735 bp in the western Mediterranean, where the low-antioxidant GWAS SNP allele was associated with lower expression (cor= –0.18, *p*=0.02), but not globally (cor<0.01). Likewise, *FRO7* expression (but not *FRO6*) was weakly associated with allelic variation in chr5: 20206030 bp in the western Mediterranean (cor= –0.19, *p*=0.01), but not globally (cor=0.07).

The *PHT4;4* antioxidant SNP covaried with our color phenotypes in a way consistent with Miyagi et al. (2015)’s findings on xanthophylls and ß-carotene. The variant associated with low antioxidant activity was significantly associated with lighter leaves and leaf extracts (i.e., higher values of PC1_LeafColor_ and PC1_ExtractColor_) in our high light (Wilcox-tests *p*=0.004), night-freezing (*p*=0.02) and no night-freezing experiments (*p*=0.007), suggesting the low-antioxidant *PHT4;4* variant is associated with low carotenoids. We did not find this association in the low light treatment (*p*=0.12), consistent with *PHT4;4* expression being upregulated under high light (Miyagi et al., 2015).

#### Physiological effects of PHT4;4 mutations

We further investigated the phenotypic effects of *PHT4;4* mutations in response to high light (vs. low light) under cold using *PHT4;4* insertion lines (CS444342, Salk_082875, and CS26443) in two different genetic backgrounds (*i.e.*, wild-type: Col-0 for the first two mutants and Landsberg erecta (LE; for the third) (Supplemental Table S3) and a subset of western Mediterranean ecotypes that segregate at the *PHT4;4* GWAS SNP (Supplemental Dataset S1).

We found that the speed of NPQ induction was significantly different among ecotypes (ANOVA, *p*<0.001), being on average 12% slower in ecotypes with the low antioxidant GWAS allele compared to ecotypes with the other allele (*p*=0.006; Figure 8 A and B). In agreement with Miyagi et al. (2015)’s results on *PHT4;4* knockouts, our results were conditioned on high light exposure: the slower NPQ induction in ecotypes with the low-antioxidant allele was only detected when plants were exposed for 2 days to high light but not when plants continuously grew under low light (Figure 8 A). Accordingly, ecotypes with the low-antioxidant GWAS allele had a 27% average reduction in *PHT4;4* expression under high light compared to ecotypes with the other allele (*p*=0.001; Figure 8 B).

**Figure 8.**
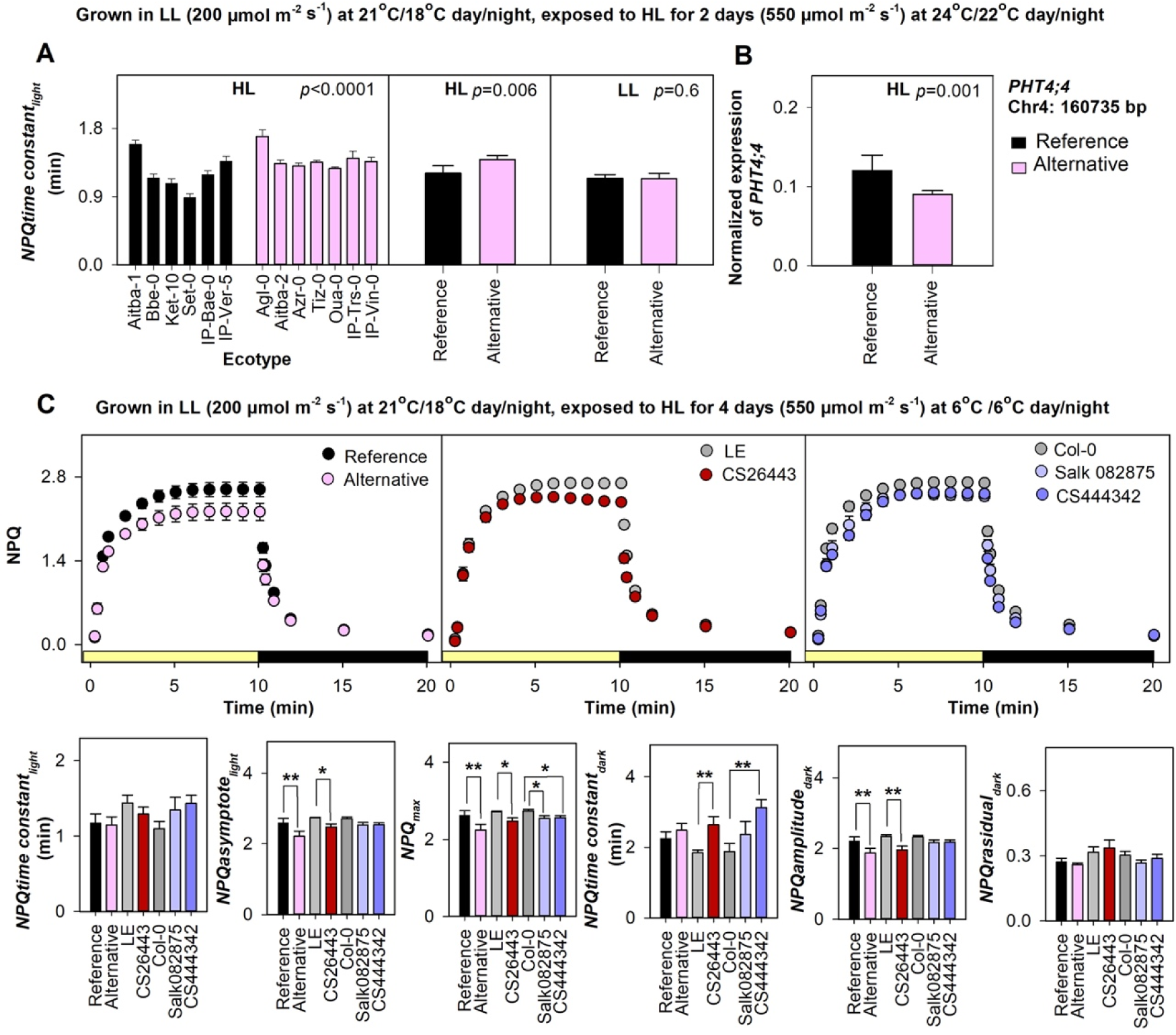
Effects of *PHT4;4* variation in NPQ kinetics and gene expression. **A** Speed of NPQ induction (*time constant_light_*; time needed to achieve 63% of the maximum capacity of NPQ) for plants grown under high light (HL: 550 μmol m^-2^ s^-1^) vs. low light (LL; 200 μmol m^-2^ s^-1^) at room temperature (5–6 replicates per ecotype), with ANOVA *p*-values indicating significant differences among ecotypes (left panel) and between ecotype groups with different GWAS SNP alleles (middle panel). **B** Significant downregulation in normalized *PHT4;4* expression in ecotypes with the GWAS SNP allele vs. ecotypes with the other allele when grown under HL (5– 6 replicates per ecotype). **C** NPQ response to light and dark cycles for plants grown under HL and cold (6°C). *PHT4;4* knockouts CS26443, Salk_082875, and CS444342, are preceded by corresponding wild-type LE and Col-0. Genotype means are depicted with standard errors based on 4–6 replicates. Asterisks indicate significant differences between ecotypes with different GWAS SNP alleles or between mutants with their corresponding wild-type based on a Dunnett’s test adjusted to multiple comparisons (*p*≤0.05).

Furthermore, a prolongation of high light exposure to 4 days combined with cold (day and night at 6°C) revealed that ecotypes with the low-antioxidant GWAS allele resembled *PHT4;4* insertion lines in their NPQ kinetics. For instance, the steady state of NPQ under light (*NPQasymptote_light_*) and the range of NPQ change under dark (*NPQamplitude_dark_*) were 15% lower (*p*=0.01) in ecotypes with the low-antioxidant allele vs. ecotypes with the other allele, and 9% (*p*=0.02) and 16% (*p*=0.008) lower, respectively, in the *PHT4;4* knockout CS26443 vs. its LE wild-type (Figure 8 C).

In a separate experiment of plants exposed for 7 days to high vs. low light (day and night at 4°C), the phenotypic effects of different *PHT4;4* variants were not as congruent between natural variation and insertion lines (Supplemental Results S3). Under high light, all western Mediterranean ecotypes had the lowest antioxidant activity, brightest/most transparent leaf extracts, and lowest chlorophyll content (Supplemental Figure S8 B–D). *PHT4;4* insertion lines showed lower chlorophyll content conditioned on high light exposure relative to their respective wild-type, consistent with our observation that in western Mediterranean ecotypes with the low-antioxidant GWAS allele, chlorophyll content was generally lowest (Figure S8). However, insertion lines and wild-type did not differ in antioxidant levels (ANOVA *p*=0.3–0.5) while neither did western Mediterranean ecotypes with different GWAS SNP alleles (ANOVA *p*=0.7).

Taken together, ecotypes with the low-antioxidant GWAS allele behaved similarly to *PHT4;4* insertion lines under high light, in that they had low *PHT4;4* gene expression, slow NPQ kinetics, and lower chlorophyll content. However, *PHT4;4* insertions did not affect total antioxidants, which were higher compared to the low antioxidant activity in our subset of western Mediterranean ecotypes. This inconsistency could be due to a *cis*-regulatory natural variant (among other explanations) of *PHT4;4* having distinct phenotypes from *PHT4;4* knockouts.

#### Potential cis-regulatory variants in the Moroccan PHT4;4 haplotype

To look for potential causal *cis-*regulatory variants, we scanned for enriched sequence motifs in the 3.5 kbp upstream and 3.5 kbp downstream region of *PHT4;4* in 63 ecotypes (61 western Mediterranean, 1 from Japan, and 1 from Czech Republic) with the low-antioxidant GWAS SNP allele/haplotype (Supplemental Figure S9 A), in contrast to 53 closely related ecotypes with the high-antioxidant allele (Supplemental Dataset 1). We discovered 54 significantly enriched motifs (p<0.05) in the low-antioxidant allele ecotypes, one of which was lacking in the high-antioxidant allele group, six of which were fixed, and 46 of which were present in more than 90% of the ecotypes with the low antioxidant allele. When we cross referenced the discovered motif sequences with the plant JASPAR (Castro-Mondragón et al., 2022) database of experimentally defined plant transcription factor binding sites (TFBS), the 54 discovered motifs in the low-antioxidant allele group had a significant overlap with 339 known TFBS. The motif lacking in the high-antioxidant allele group overlapped with four known TFBS in the HD-ZIP family (MA0990.1, MA1212.1, MA1369.1, MA1375.1). Members of this family function in regulating stress response and plant growth (Sharif et al., 2021). Amongst the most common TFBS across the discovered motifs were members of the DOF family, with DOF1.7, DOF5.8 and DOF6.3 being present in 20%, 18.5% and 15.4% of the total discovered motifs respectively (Table 2, Supplemental Figure S9 B, C). Members of this family function in regulating growth, development, and biotic and abiotic stress response (Zou and Sun, 2023). Lastly, motifs in the MIKC TFBS family were common in more than 10% of the discovered motifs (Table 2). Members of this family function in regulating cyclical developmental processes in response to cold (Castelán-Muñoz et al., 2019).

**Table 2.** Most frequent transcription factor binding sites (TFBS) found within 54 discovered motifs 3 kbp around *PHT4;4* in 63 western Mediterranean lines carrying the low-antioxidant *PHT4;4* haplotype. For each known TFBS, we report the number of overlapping motifs (N).

| TFBS id | TFBS name | TFBS family | N motifs |
| --- | --- | --- | --- |
| MA1277.1 | DOF1.7 | DOF | 13 |
| MA1267.1 | DOF5.8 | DOF | 12 |
| MA1274.1 | DOF6.3 | DOF | 10 |
| MA1184.1 | RVE1 | Myb-related | 9 |
| MA1191.1 | RVE7L | Myb-related | 9 |
| MA1268.1 | CDF5 | DOF | 9 |
| MA1273.1 | DOF4.2 | DOF | 9 |
| MA1279.1 | DOF1.5 | DOF | 9 |
| MA0563.1 | SEP3 | MIKC | 8 |
| MA1012.1 | AGL27 | MIKC | 7 |
| MA1204.1 | AGL13 | MIKC | 7 |
| MA1205.1 | AGL6 | MIKC | 7 |
| MA1281.1 | DOF5.1 | DOF | 7 |

## Discussion

Arabidopsis exhibits natural genetic variation in abiotic stress response, which may be linked to phenological and physiological adaptation along elevational gradients. Germination and flowering time show evidence of local adaptation in response to seasonal cold at high elevations and to early drought at low elevations (Méndez-Vigo et al., 2011; Suter et al., 2014; Vidigal et al., 2016). Water use efficiency (WUE) and its genetic basis may be associated with phenology (Kenney et al., 2014; Lovell et al., 2013), though changes in WUE with elevation have been less studied (DeLeo et al., 2020; Des Marais et al., 2014; Dittberner et al., 2018; Mckay et al., 2003; Wolfe and Tonsor, 2014). Elevational clines related to dwarfism evolution at high elevations have been identified (Luo et al., 2015a, b; Singh and Roy, 2017; Tripathi et al., 2019; Tyagi et al., 2016). However, past studies have typically focused on individual geographic regions in Eurasia, precluding a broader understanding of adaptation to global elevational gradients.

Here we examined the range-wide genomics and physiology of adaptation to elevation of Arabidopsis using a multi-regional approach informed by phylogeography. We uncovered elevational clines indicating changing selection for several traits and genomic variants. Most clines were region-specific, despite similarity between regions in elevational environmental gradients, where differences in phenology are likely part of a coordinated phenology-physiology strategy that differs between regions. Antioxidant activity, pigmentation, and NPQ kinetics showed substantial genetic variation. These traits may be particularly important to local adaptation to Moroccan mountains. We showed that even when selective environmental gradients are consistent among mountain ranges, patterns of multivariate adaptation to elevation and their genetic basis can be unique to each mountain range (see Tusiime et al., 2020; Wos, et al. 2022). This could be due to phenotypic and genetic heterogeneity (sensu López-Arboleda et al., 2021) across different regions affected by subtle differences in selective gradients and phenotypic clines despite similar environmental clines, or by the historical contingency of each region. Our regional approach in a widespread model plant highlights that a better integration of plant adaptation at a range-wide scale will help improve evolutionary predictions (*e.g.*, in globally important crops or in invasive species) under the current scenario of accelerated anthropogenic global change.

### Coordinated life history and physiology along elevation

In Asia, high elevation ecotypes showed a putatively adaptive strategy of rapid and plastic phenology (*i.e.*, more rapid at 16°C than at 10°C) and low water and nitrogen use. High elevation genotypes with low δ^13^C tended to harbor mutations in *FLM*, a flowering time regulator, suggesting a potential pleiotropic effect of flowering time genetic control on water use efficiency in Asia. Previous studies have found genotypic correlations between low WUE and earlier flowering both due to variation across the genome (Kenney et al., 2014) and specifically to pleiotropic effects of a flowering time regulator (*FRI*) (Lovell et al., 2013). Furthermore, studies in the western Himalayas suggest that increasing cold and light intensity with elevation influences life history and genomic variation (Singh and Roy, 2017; Tripathi et al., 2019; Tyagi et al., 2016). Here, two ecotypes from high elevation central Asia showed a strong increase of NPQ kinetics after cold exposure. High light and low pCO_2_ induces drought stress in leaves via ABA signaling and stomatal dynamics (Kuromori et al., 2022; Moustakas et al., 2022), while high light and cold increases the risk of photo-oxidation (Wise, 1995). We hypothesize that in Asia high elevation ecotypes exhibit a multivariate stress-escape strategy involving rapid NPQ and a fast ‘spring annual’ life cycle.

In the western Mediterranean, high elevation ecotypes showed a putatively adaptive strategy of delayed flowering and high water and nitrogen use efficiency, suggesting drought avoidance. By contrast, low elevation ecotypes had open rosettes associated with a rapid-cycling strategy (early flowering) potentially allowing drought escape (Ludlow, 1989). The genetic basis of this cline is likely partly caused by allelic variation in the seed dormancy/flowering time *DOG1* gene (Graeber et al., 2014; Huo et al., 2016), and in two candidate genes related to abiotic stress response in the ABA pathway, *MAP3k* and *NIC3* (Kim et al., 2019; Takahashi et al., 2020). Previous studies in the Pyrenees have shown that earlier flowering relates to drought-escape in the lowlands (MontesinosLJNavarro et al., 2009, 2011; Wolfe and Tonsor, 2014), and our results indicate that this strategy may involve evolution of ABA signaling. Ecotypes from high elevation have alleles that allow them to counteract water loss under stress, which in turn is adjusted with a ‘winter-annual’ life cycle, but not those from low elevations which are probably less commonly exposed to environmental stress because of their rapid cycle during wet periods (Picó 2012).

### Traits associated with protection from photo-oxidation

Ecotypes showed shared increases in antioxidants and changes in leaf color in response to bright light and night-time freezing, but little associations with elevation of origin. Photo-oxidation is also a threat at lower elevations due to drought, potentially resulting in no clear change in selection (Nakabayashi et al., 2014). Furthermore, while changes in antioxidants and pigments mediate high light acclimation (Aarabi et al., 2023), they might not indicate cold acclimation (Distelbarth et al., 2013), but could be related to freezing tolerance via flavonoid accumulation (Hannah et al., 2016). The combination of cold and high light can thus comprise diverse metabolic and genomic pathways, that could be region-specific (see next section).

Afroalpine populations occupy an outlier environment with respect to clear sky radiation and night-time cold throughout the year. Here, antioxidant activity appeared to increase with elevation under night-time freezing. Afroalpine ecotypes also had yellower leaf extracts compared to dark orange and deep red extracts in other regions, potentially indicating less xanthophylls used in NPQ, a hypothetical reduced sensitivity to cold (Havaux and Kloppstech, 2001). More orange extracts and higher plasticity in extracts color were associated with higher survival under night-time freezing, suggesting potential selection on pigment content. Additionally, a high elevation Afroalpine ecotype was very small, had little total carotenoid content, with slower but greater NPQ under cold, and distinctly high leaf dry matter content under controlled conditions. These characteristics could be adaptive if they enhance freezing resistance (Gorsuch et al., 2010).

We found significant genetic variation in NPQ kinetics parameters in response to cold and drought (i.e., ecotypes were significantly different in their response), but this variation was not clearly related to elevation of origin. Cold resulted in slower NPQ induction and relaxation across genotypes, while drought resulted in highly variable NPQ induction and relaxation among ecotypes. NPQ can remove over 75% of absorbed light energy, thus its fast induction in response to stress can play an essential role for rapid defense against damage from photoinhibition (Malnoë, 2018). Slow relaxation of NPQ (particularly evident in two Asian highland ecotypes) could limit NPQ and plant growth, but provide better protection to cold (Kromdijk et al., 2016). We hypothesize that the generally slower NPQ speed in cold vs. drought is due to temperature-dependent changes in the activity of both NPQ-related enzymes, *violaxanthin de*-epoxidase and *zeaxanthin* epoxidase (Nilkens et al., 2010).

### Natural and knockout *PHT4;4* variants differ in NPQ induction and gene expression under high light

Under bright and cool conditions, western Mediterranean ecotypes (mostly from Morocco) had the lowest antioxidant activity, chlorophyll content, and slow NPQ kinetics. These ecotypes have almost exclusively alleles near a transporter gene for ascorbate into the chloroplast (*PHT4;4*) associated with low antioxidant activity under high light, and from the subset we tested under high light, they showed a significant downregulation of *PHT4;4* expression relative to ecotypes with the other allele. Previous studies (Miyaji et al., 2015) showed that the level of *PHT4;4* expression increases with light exposure. *PHT4;4* knockouts under high light had slower NPQ induction, lower levels of reduced ascorbate in leaves, and lower levels of xanthophyll and beta-carotene (Miyaji et al., 2015). Here we show that under high light and cold, *PHT4;4* knockouts and western Mediterranean ecotypes with the low-antioxidant GWAS allele had lower NPQ and chlorophyll content. Ascorbate is an antioxidant and cofactor used in the xanthophyll cycle which in turn aids induction of NPQ. The western Mediterranean ecotypes carrying the distinct *PHT4;4* haplotype linked to our GWAS SNP, might be tolerating high light and cold conditions by maintaining a low but steady NPQ speed involving downregulated *PHT4;*4, while reducing oxidative risk by producing less chlorophyll by, for example, reducing light-harvesting antenna pigments of Photosystem II (Havaux and Kloppstech, 2001), which could incur a cost under lower light conditions. This strategy might allow to limit photo-oxidative damage independently from antioxidant activity in the bright and seasonally cold Atlas Mountains.

Furthermore, we showed that the western Mediterranean low-antioxidant GWAS *PHT4;4* haplotype (compared to related ecotypes lacking this haplotype) has putative *cis*-regulatory motifs enriched primarily for transcription factor binding sites (TFBS) in the DOF zinc-finger family. This family is involved in hormonal response and photoperiodic regulation during plant development and under stress (Zou and Suri, 2023). DOF response elements are present in the promoter region of several cold-induced genes which expression is exclusively upregulated in response to light exposure under cold (Soitamo et al., 2008). We also identified motifs associated with TFBS in the MIKC family, which are involved in floral development and are key components of gene regulatory networks involved in stress response and developmental-phenological plasticity in response to seasonal cold (Castelán-Muñoz et al., 2019). Furthermore, our analysis revealed a motif only present in the western Mediterranean low-antioxidant GWAS *PHT4;4* haplotype, that overlapped with four known TFBS in the HD-ZIP family. Genes involved with this family are critical for plant growth and development under stressful environmental conditions (Sharif et al., 2021). We believe the evidence may indicate a *cis*-regulatory variant at *PHT4;4* that reduces its expression under high light causing slower NPQ kinetics. This phenotype might be part of a more conservative growth strategy that does not require rapid NPQ kinetics, possibly due to lower chlorophyll levels in ecotypes from the region.

Finally, *PHT4;4* knockouts might not show antioxidant phenotypes due to epistasis with other genes in western Mediterranean ecotypes, like the top GWAS QTL *FRO6/7*, an iron transporter with similar allele frequency to the *PHT4;4* SNP. Nam et al. (2021) found prevention of downregulation of photosynthesis related genes via ascorbate accumulation by *PHT4;4* induced by transcription factors responsive to iron deficiency. We hypothesize that the interaction between *PHT4;4* and *FRO6/7* transcription factors in response to high light and cold could influence the low antioxidant and chlorophyll content of western Mediterranean ecotypes with downregulated *PHT4;4*. Indeed further validation studies with mapping populations or transgenics targeting the specific natural variation found in the western Mediterranean are needed. Our findings highlight the complexity of investigating the underpinnings of the functional genomic and phenotypic variation in natural systems.

## Materials and Methods

### Plant material

We selected a diverse set of 271 ecotypes (i.e., naturally inbred genotypes; Supplemental Dataset S1) which we collectively call our diversity panel: 232 from the 1001 Genomes Project (ABRC stocks), 21 from Morocco and Tanzania (NASC stocks), and 18 newly collected lines (8 of them here sequenced; Supplemental Methods S1) along elevational gradients in Ethiopia (in 2017) and Uganda (in 2018) ranging from 3691–4374 m. We specifically targeted several regional elevational gradients with this diversity panel, which we subsequently classified into **five regions** following genetic cluster (according to Alonso-Blanco et al., 2016 and Durvasula et al., 2017, see Supplemental Figure S1) and geographic origin:

**1)** Asia (46 ecotypes; 45 in the Asia genetic cluster + 1 admixed from Kazakhstan).
**2)** central Europe/Caucasus (59 ecotypes; 16 in the Central Europe cluster + 37 in the Italy-Balkan-Caucasus cluster + 6 admixed from Germany, Italy, Lebanon, Switzerland, and Turkey).
**3)** northwestern Europe (26 ecotypes; 6 in the Germany genetic cluster + 3 in the North Sweden cluster + 9 in the South Sweden cluster + 4 in the Western Europe cluster + 4 admixed from the Netherlands and the United Kingdom).
**4)** western Mediterranean (112 ecotypes; all from Iberia and Morocco: 13 in the Relict genetic cluster + 53 in the Spain cluster + 9 in the Western Europe cluster + 22 in the Moroccan clusters + 15 admixed from Spain).
**5)** eastern Africa (19 ecotypes, only 9 sequenced; 16 from Ethiopia + 2 from Uganda + 1 from Tanzania).

The remaining accessions were from North America and Japan, thus designated as unassigned, along with one accession from Cape Verde given how different this location is from all other regions (9 ecotypes). With subsets of our diversity panel, we conducted three large-scale growth chamber experiments (Experiment 1–3 below, Supplemental Dataset S1), and a detailed study on NPQ kinetics in response to cold and drought. When trait measurements (explained below) could be averaged across biological replicates, we obtained breeding values of phenotypes from the fixed effects of linear mixed models that estimated the trait mean per ecotype with ‘tray’ as a random factor in the R lme4 package. All phenotypes were measured in adult pre-flowering plants.

### Seed bulking and initial growing conditions

Seeds from 271 ecotypes were bulked after dark-stratification in distilled water for 4 d at 4°C, in a growth chamber at 25°C day/15°C night with a 16 h day/8 h night photoperiod in a reach-in growth chamber (Model PGC-40L2, Percival Scientific, IA, USA). One 21 d old plant per pot was kept and vernalized for 10 wk. at 4°C with an 8 h day/16 h night photoperiod, then returned to initial growth conditions until harvesting. Harvested seeds were dark-stratified as in the bulking before performing experiments in the Percival, then grown in cone-tainers (1.5-inch diameter, 164 ml) in a RL98 rack with nutrient rich soil (Supplemental Methods S2). After the first true leaves expanded (ca. 23 d old plants), pots were thinned to one plant per pot. We used a 12 h day/night photoperiod for all large-scale experiments, randomized pots withing trays, and periodically rotated trays and moved them around the growth chamber (or within shelves) to prevent positional effects.

### Experiment 1: Low pCO_2_ in 253 ecotypes

We randomly selected 140 ecotypes for 5 biological replicates (individual plants) and 113 ecotypes for 4 biological replicates, for a total of 1152 plants. The difference in number of replicates was due to space constraints in the growth chamber. We imposed a treatment of low pCO_2_ by scrubbing CO_2_ from the Percival air using a soda lime absorbent material (SODASORB HP, Divers Supply Inc., LA, USA). We set the growth chamber to 200 ppm CO_2_, which at our elevation of ∼350 m corresponds to ∼19.5 Pa, the approximate pre-industrial partial pressure of CO_2_ at 3000 m, compared to a pre-industrial partial pressure of ∼28 Pa at sea level. We sought to mimic high elevation conditions by growing plants under high light (∼462–616 μmol m^−2^ s^−1^) and cool conditions (∼3–13°C; Supplemental Dataset S3) with moderate soil drying. We terminated the experiment 156 d after planting and subsequently completed harvesting and phenotyping within 5 d.

### Phenotypes in Experiment 1

We recorded phenotypic variation in six traits on 250–253 ecotypes: δ^13^C, δ^15^N, root diameter, rosette diameter, and rosette compactness. Just prior to harvest, we photographed overhead rosettes of each plant and used ImageJ (Schneider et al., 2012) to measure rosette diameter (as the average between the longest and the shortest distance between opposite leaf tips) and rosette compactness (by assigning a rating from 1–5 where 5 corresponded to more compact rosettes with shorter petioles and total leaf overlap, thus less soil exposure and 1 corresponded to less compact rosettes with longer petioles and minimal leaf overlap, thus more soil exposure). After harvesting, root diameter for each plant was measured 3 mm below the hypocotyl/root junction with a caliper. We then used 2–4 young healthy leaves per plant for estimating δ^13^C and δ^15^N. Leaves of each ecotype were pooled, dried at 27°C for 24 h, and homogenized to a fine powder. We sent 1.5–2.5 mg of fine powder per ecotype to the University of California Davis Stable Isotope Facility where δ^13^C and δ^15^N were quantified following international standards using an elemental analyzer interfaced to a continuous flow isotope ratio mass spectrometer (Sharp, 2005). We measured stomatal density on a subset of 80 widely distributed ecotypes (Supplemental Dataset S1): 2–4 young healthy leaves were collected from 2–3 plants per ecotype and stored in 75% ethanol at 4°C. Epidermal peels from the abaxial surface of each leaf were softened in distilled water for 3–7 min, immediately placed into 0.02% Toluidine Blue stain for 3–5 min, rinsed with distilled water for 3 s, and put onto a slide with a small drop of distilled water for imaging. Images were taken at 20X magnification using a Zeiss compound microscope and stomatal density (stomata/mm^2^) was quantified in ImageJ, using distance-calibrated images of one representative area of each epidermal peel. To assess coordinated life history variation among regions, we summarized variation in phenotypes from Experiment 1 and days to flower (in plants grown at 10 and 16 °C; published data in Alonso-Blanco et al. 2016) with a principal component analysis (PCA; R ‘prcomp’ function).

### Experiment 2: Low vs. High Light in 111–114 ecotypes

We conducted Experiment 2 using identical growth parameters as above except for light intensity (Supplemental Dataset S4). Five biological replicates of 111–114 ecotypes, for a total of 1125 plants, were grown under two light levels under cool conditions (∼11°C day/3°C night). Plants grown on the lower level of the growth chamber received the “high light” treatment with 100% maximum light output (∼616 μmol m^−2^ s^−1^), and plants grown on the upper level of the chamber received the “low light” treatment with 57% maximum light output (∼350 μmol m^−2^ s^−1^). We distributed plants this way, as the upper level of the chamber tends to get warmer, balancing out the heat from bright lights in the bottom shelf. Soil temperature after 43 d taken with a FLIR ONE (Gen 2) thermal camera for iOS (FLIR Systems, USA) showed that differences in soil temperature were on average only within 1.3°C, with warmer pots under the high light treatment (12.4°C vs. 11.1°C).

### Experiment 3: Night-time above vs. night-time below freezing in 170–172 ecotypes

Experiment 3 consisted of two consecutive experiments using identical growth parameters as Experiment 1, except for night-time temperatures (Supplemental Dataset S5). Six replicates of 170–172 ecotypes were grown under night-time below freezing temperature (850 plants) and consecutively under night-time above freezing temperature (860 plants). Both experiments maintained high light (∼462–616 μmol m^−2^ s^−1^) and cool conditions throughout (∼15°C day/4°C night), but in the night-time freezing experiment we introduced below freezing temperatures (from –4 to –2°C) after 40 d for 30 d. We monitored soil temperature with probes every 30 min, which did not reveal soil frost during the day.

### Phenotypes in Experiment 2 and 3

We recorded phenotypic variation in three traits related to photosynthesis: fresh aboveground biomass, antioxidant activity, and leaf color. Ecotypes were harvested 82–89 d (Experiment 2) and 72–75 d (Experiment 3) after planting. Biological replicates from each ecotype were pooled together (due to their small size) and cleaned with distilled water for assessing collective fresh aboveground mass and antioxidant activity with 1,1-diphenyl-2-picrylhydrazyl (DPPH), an assay that estimates free radical scavenging activity of antioxidants (protocol in Seyoum et al., 2006). Leaf color was estimated with ImageJ, excluding late germinants. In Experiment 2, rosette leaf color was measured from the pixels selected at a representative location within a mid-aged leaf on rosette photographs of each plant taken just prior to harvest. We used the option “Color Histogram” to obtain the variation and calculate percent of red, green, and blue pixels (RGB values). In Experiment 3, we obtained variation in RGB values from images of the pooled leaves extracts per ecotype in test tubes used for DPPH. Images were white-balanced and zoomed into the test tube avoiding any discolored, dark, or bright areas. We note that upon visual examination of photographs, rosette colors are probably more indicative of anthocyanins, while extract colors may also inform on other pigments like carotenoids, including xanthophylls. We then used PCA to summarize variation in color assessed from RGB values in contrasting conditions. Based on the color space in PC1 and PC2, we calculated plasticity in color: the Euclidean distance between ecotype values of the two contrasting conditions using the Pythagorean theorem. We also calculated plasticity in antioxidant activity by finding the difference between experimental conditions (stressful minus control condition). Survival rates per ecotype were obtained from the number of replicates that survived at the end of each experiment divided by the number of replicates that germinated within treatments. We used survival as a proxy of fitness and analyzed how other traits affect it with Pearson correlations.

### Assessing region-specific climatic and trait variation

Based on ecotypes with accurate coordinates and elevation of origin, we examined how study regions varied climatically and differed in phenotypes. We first evaluated regional variation of environmental variables on elevation using linear models (R ‘lm’ function) and performed a PCA on 85 environmental variables (Supplemental Dataset S2). We then used ANOVA to test for differences in trait variation among regions, followed by Tukey’s honest significant difference post-hoc tests to detect pairwise regional differences. We also used ANOVA to test for significant differences in trait variation between conditions (plasticity) in Experiments 2 and 3.

### Assessing region-specific putatively adaptive trait-elevation clines

For each trait of interest, including the first two PCs of Experiment 1–3 described above, we tested for associations with elevation within regions using linear mixed effects kinship models with the function ‘lmekin’ in the kinship2 R package (Therneau, 2012). These models accounted for population genetic structure by incorporating a kinship matrix of ecotypes as a random effect. When significant, they thus provide evidence of selection maintaining trait-elevation clines. A kinship matrix of sequenced ecotypes was obtained from a published VCF of genotypes (Durvasula et al., 2017), which we concatenated with the here sequenced eastern Africa ecotypes (Supplemental Methods 1), for a total of 1233 ecotypes (261 in our study) in that VCF. First, this VCF was converted to gds format for manipulation in the SNPRelate R package (Zheng et al., 2012). We then visualized population genetic structure with a neighbor-joining tree with a subset of 562,194 unlinked SNPs obtained with ‘snpgdsLDpruning’ in SNPRelate with a linkage-disequilibrium threshold of 0.5 and a 10 kb window. With this set of SNPs, we calculated a distance matrix based on genetic dissimilarity between ecotypes and used the ‘nj’ and ‘plotnj’ function in the R package ape (Paradis et al., 2004). We obtained a kinship matrix with the snpgdsIBS function (*i.e.*, based on the fraction of identity by state for each pair of samples) and the same set of unlinked SNPs.

### Assessing genome-wide phenotypic associations and genomic clines

We imputed the above VCF based on LD with Beagle v.4.1 (Browning et al., 2018), resulting in 22,619,232 SNPs of which 11,500,298 were polymorphic. We used univariate linear mixed effects models in gemma v.0.98.3 (Zhou and Stephens, 2012) to perform genome wide association studies (GWAS). Because traits generally differed in the number of analyzed individuals, we filtered out loci with maf<0.05 directly in gemma. To account for multiple testing, we used a false-discovery-rate (FDR) threshold of 0.05 to detect significantly associated SNPs with traits of interest. We ran gemma on the global dataset for all traits, as well as by region for those traits that showed a significant association with elevation. For flowering time regional GWAS, we used all the ecotypes with available data: 60 for Asia, 289 for central Europe/Caucasus, 398 for northwestern Europe, and 183 for western Mediterranean. We identified QTL by assigning the top SNPs (i.e., those with the lowest 100 p-values for each trait) to candidate genes using the Araport11 reannotation (Cheng et al., 2017) and a 5 kb window. To examine if selection on elevation maintained allelic variation in top SNPs in putative QTL, we evaluated the association of allelic variation with elevation while accounting for population genetic structure with ‘lmekin’.

### Assessing NPQ kinetics in a global panel of 9–11 ecotypes

We performed a detailed study on NPQ kinetics based on ecotypes with a global distribution representing the lowest (22–254 m) and highest (1576–4078 m) elevational range of Arabidopsis, that also showed great variation in antioxidant activity. With this panel we aimed to capture diverse strategies, but given the limited number of ecotypes, not necessarily make conclusive cline tests.

Seeds were dark-stratified at 4°C for 4 d in water (or in 800 PPM gibberellic acid for the JL011-2-1 ecotype), sown in 8.89 × 8.89 cm pots with soil-less BM2 potting mix under moist conditions, and grown in a reach-in Percival chamber (model AR-66L2). Growing conditions were set at 21°C day/18°C night with an 8 h day/16 h night photoperiod (200 μmol m^−2^ s^−1^) and 60% RH, 5 d old plants thinned to one per pot, well-watered, and trays repositioned at random locations in the chamber.

Two weeks after germination, NPQ kinetics was measured on seedlings in two separate experiments (5–10 biological replicates per ecotype per treatment) using a fluorescence imager (Closed FluorCam 7 FC 800-C, Photon System Instruments, Czech Republic). Half of the seedlings were subjected to drought (5 d of no watering; 2 ecotypes excluded due to flowering) or to 17 h of overnight chilling at 6°C, and the other half were maintained under normal conditions and used as control. Plants were subjected to three cycles of 3 min 1000 μmol m^-2^ s^-1^ followed by 2 min of dark. NPQ of chlorophyll fluorescence was determined assuming the Stern-Volmer quenching model (NPQ = *F*_m_/*F*_m_′ – 1; Bilger and Bjorkman, 1994). NPQ was then calculated at following intervals of 10s, 10s, 20s, 20s, 60s and 60s in light and 10s, 20s, 30s and 60s in dark. To this end, raw images were processed manually using a FluorCam 7 by drawing three circles per plant of 170–180 pixels on three young fully developed leaves. The NPQ data were fitted to exponential equations to parametrize the NPQ kinetics of its induction in light and relaxation in dark of three subsequent light-dark cycles (Sahay et al., 2023, Supplemental Figure S3) using a custom-made script in MATLAB v. R2018b. After 30 d since germination, we harvested rosettes from control conditions to estimate fresh and dry aboveground biomass, and photosynthesis related pigments. For each ecotype, we harvested 4–6 biological replicates, which were immediately weighted to obtain fresh weight. To estimate dry weight, fresh rosettes were dried to constant weight at 70°C. On the remaining 4–6 biological replicates, we quantified photosynthesis related pigments from 2.32 cm^2^ leaf tissue from the interveinal region following a 100% methanol-based extraction (Lichtenthaler, 1987). To understand variation among ecotypes in NPQ kinetics, biomass, and pigments, we performed ANOVA and Tukey HSD post-hoc tests (α=0.05) in SAS v.9.4.

### Assessing the physiological effect of allelic variation and gene expression of *PHT4;4* in western Mediterranean ecotypes

We investigated the effect of allelic variation in *PHT4;4* (chr4: 160735) on western Mediterranean ecotypes, using seven ecotypes with the low-antioxidant allele and six closely related ecotypes with the high-antioxidant allele (Supplemental Dataset S1), along with three *PHT4;4* knockouts. We obtained a Ds transposon tagged mutant in a Ler genetic background (CS26443) and two T-DNA insertional mutants in a Col-0 genetic background (Salk_082875 and CS444342) from the Cold Spring Harbor Laboratory and the Arabidopsis Biological Resource Center (Ohio State University), respectively. Lines were selfed until a homozygous progeny for knockouts was confirmed by PCR (Supplemental Table S3). Two weeks after germination under control conditions, half of the seedlings were subjected to two days of high light (HL; 550 μmol m^−2^ s^−1^ at 24°C/22°C day /night with an 8 h day/16 h night photoperiod) or to four days of HL under cold (550 μmol m^−2^ s^−1^ at 6°C with a 16 h day/8 h night photoperiod) in a walk-in growth chamber (Conviron model GR48, Controlled Environments, Manitoba, Canada). The other half were maintained under normal conditions and used as controls. NPQ kinetics was investigated during 10 min of light and 10 min of dark, and parametrized as in the global NPQ experiment, except that saturating flashes were spaced at the following intervals (in s): 15, 30, 30, 60, 60, 60, 60, 60, 60, 60, 60, 60, 60, 9, 15, 30, 60,180, and 300.

To quantify *PHT4;4* expression under high light, four leaf discs (2.32 cm^2^) from a fully expanded leaf of 5–6 biological replicates were collected 2 h after the photoperiod started in ecotypes treated for 2 days with HL at RT (first HL experiment above). Total mRNA was extracted from leaf tissue with a NucleoSpin RNA/Protein kit (REF740933, Macherey-Nagel GmbH & Co.), then treated with DNase before transcription into cDNA using a Superscript III First-Strand Synthesis System (18080-051, Thermo Fisher Scientific). The cDNA was amplified by quantitative reverse transcription PCR (RT-qPCR) to quantify the *PHT4;4* transcript (5’-GCATATAGCTCTCCCAAGGATG-3’ and 5’-TAGAGACCCGACTGAGAGAATG-3’) relative to the elongation factor (AtEF1α; 5’-TGAGCACGCTCTTCTTGCTTTCA-3’ and 5’-GGTGGTGGCATCCATCTTGTTACA-3’). The RT-qPCR was performed using a SsoAdvanced Universal SYBR Green Supermix (172-5270, BioRad). We used ANOVA followed by a Dunnett’s test to investigate differences in physiological traits and *PHT4;4* gene expression between ecotypes with different GWAS alleles and between knockouts and corresponding wild-type.

We performed an additional experiment to quantify antioxidant activity in *PHT4;4* knockouts and their respective wild-type, along with five western Mediterranean ecotypes that differ in the GWAS allele (Supplemental Dataset S1). Seeds were dark stratified at 4°C in gibberellic acid (1000 PPM) for 7 d, then sown in 4 × 4 cm pots with BMX potting mix and kept moist. We grew plants for 20 d in a walk-in chamber (Conviron model CMP4030, Controlled Environments, ND, USA) with a 16 h day/8 h night photoperiod at ∼22°C day/18°C night, and subsequently transferred them to a cold room at 4–8°C day and night, where 10–16 randomized biological replicates of each genotype were placed under high (∼600 μmol m^−2^ s^−1^) or low light (∼150 μmol m^−2^ s^−1^). We started with an 8 h day/ 16 h night photoperiod, and gradually increased it to 12 h over the course of 1 wk. After 10 d, 2 h after the photoperiod started, we measured chlorophyll content with a SPAD meter (Spectrum Technologies, IL, USA) based on an average of three points along a fully expanded young leaf, then harvested rosettes to perform the same DPPH assay as in our GWAS high light experiment. We took photographs from leaf extracts and quantified RGB colors and brightness with ImageJ in white-balanced photos. We then performed a PCA on RGB values across genotypes and used PC1 and PC2 as color traits.

### Assessing potential *cis*-regulatory variation of *PHT4;4* in Moroccan ecotypes

We used samtools v1.10 (Li et al. 2009) to extract the 3.5kbp upstream and downstream regions of *PHT4;4*. Specifically, we extracted the 3kbp upstream and 0.5kbp downstream of the gene’s start codon for the upstream regulatory coding region, and the 0.5kbp upstream and 3kbp downstream of the gene’s end codon for the downstream regulatory coding region. We then used the standard functions of the program STREME (Bailey, 2021) to calculate significantly enriched motifs within 63 western Mediterranean ecotypes carrying the low-antioxidant GWAS haplotype (Supplemental Figure S9) vs. 53 closely related ones with the other allele (based on genome-wide genetic distance, see the tree in Supplemental Figure S1). We then searched for overlaps between the discovered enriched motifs and the JASPAR 2022 CORE (Castro-Mondragón et al., 2022) plant transcription factor binding sites database with MEME suite’s online tool TOMTOM (Gupta et al., 2007).

## Accession numbers

Sequence data (bam files), a VCF with newly sequenced eastern Africa ecotypes, and phenotypes from this article are available in the Figshare public repository with DOI: 10.6084/m9.figshare.23960727

## Acknowledgments

Permits: Plant material was exported from Uganda with permission of The Ministry of Agriculture, Animal Industry and Fisheries’ Plant Quarantine and Inspection Services, permit UQIS 4414/93/PC (E). Material was exported from Ethiopia with permission of the Ethiopian Biodiversity Institute, Ref. no. EBE71/7065/2018. Material was imported to the USA under USDA APHIS permits P37-17-01651 and P37-18-00230. Chloee McLaughlin, Jeremy Sutherland, Tim Gilpatrick, Connor Campana, Victoria Meagher, and Jaden Hill assisted in experiment setup and rosette harvests. Shawn Berghard advised in pest and disease control. Alexander Batelaan, Darla Brennan, Rachel Gerdes, Bailey McLean, and Annie Nelson helped in performing NPQ kinetics assays. Funding was provided by NSF DEB-1927009 and NIH 1R35GM138300-01 awards to JRL, and by NSF IOS-2142993 to KG.

## Author contributions

DG, CL, SS, KG, and JRL designed the research; DG, CL, AH, SS, LL, TX, MT, EK, DE, JK, MY, CEB, and TW performed research; DG, CL, SS, LL, TX, MT, EK analyzed data; DG, CL, SS, KG, and JRL wrote the paper with contributions of all other authors.

## List of Supplemental Data

### Supplemental Methods

S1. Sequencing and Variant Calling.

S2. Analysis of RGB color data.

### Supplemental Results

S1. Analysis of RGB color data.

S2. Flowering time QTL in northwestern Europe.

S3. Physiological effects under high light of *PHT4;4* variants.

### Supplemental Figures

S1. NJ tree of the global population genetic structure of Arabidopsis with 1233 ecotypes.

S2. NJ gene trees of vernalization-pathway gene sequences.

S3. Representative NPQ kinetics curve and parameters.

S4. Cold effect on NPQ kinetics for three subsequent light-dark cycles in 11 ecotypes.

S5. Drought effect on NPQ kinetics for three subsequent light-dark cycles in 9 ecotypes.

S6. Aboveground biomass of 11 ecotypes grown under control conditions.

S7. Regional *DOG1* haplotype variation relative to elevation and flowering time.

S8. Antioxidant-related phenotypes under HL vs LL (cold) in *PHT4;4* variants.

S9. NJ tree and motifs linked to the low antioxidant *PHT4;4* haplotype.

### Supplemental Tables

S1. ANOVA on NPQ kinetics in response to cold and drought.

S2. Quantification of photosynthesis related pigments in 8 ecotypes. S3. Information of insertional mutants in *PHT4;4*.

### Supplemental Datasets

S1. Coordinates, elevation, and region of ecotypes used in all experiments.

S2. List of environmental variables, source, and definition used in the PCA of Figure 1.

S3–S5. Percival programs and HOBO data for conditions in Experiments 1–3.

S6. Dataset, top SNPs, and functional gene annotation in Flowering Time GWAS.

S7. Top SNPs and gene annotation for the low pCO_2_ GWAS.

S8. Top SNPs and gene annotation for the HL vs. LL GWAS.

S9. Top SNPs and gene annotation for the night-freezing vs. no freezing GWAS.

## Notes

### Competing Interest Statement

The authors have declared no competing interest.

### Summary of Updates

Includes new experiments on the physiological effects of PHT4;4 natural variants and knockouts.

