## Supplemental Data for "The genomics and physiology of abiotic stressors associated with global elevation gradients in *Arabidopsis thaliana*"

**Supplemental Methods**

**S1. DNA extraction, sequencing, and variant calling of Afroalpine ecotypes.** Total genomic DNA of 16 newly collected accessions (8 phenotyped in this study; JL samples that have a sample id in Supplemental Dataset S1) was extracted from young leaves. Prior to extraction, tissue was homogenized using a TissueLyser (Qiagen). Each sample followed two cycles of homogenization, the first cycle was done for 30 s and the second was 15 s to ensure all tissue had been pulverized. Then, the DNA was extracted using the PTB protocol (Wales and Kislter, 2019). Modifications were made to the length of the digestion (24 hours and the amount of the elution buffer (25 µl). At least 4 µg of genomic DNA was used to construct pair-end sequencing libraries (PerkinElmer NEXTFLEX Rapid XP DNA-Seq Kit HT) which were sequenced on a NovaSeq 6000 S4 X platform (Illumina, CA, USA). On average, a total of 1.22 Gb data for each accession was used for further analysis.

After trimming Illumina tags with Trimmomatic using default parameters (Bolger et al., 2014), pair-end reads from each accession were mapped to the *Arabidopsis thaliana* TAIR10 genome (Araport11 assembly) using bwa (Li and Durbin, 2009) (Version: 0.7.12) with the following parameters: bwa -n 0.01 -o 2 -l 16500 -t 7. Read alignments were converted into the BAM format, sorted according to mapping coordinates, and PCR duplicates removed using the Picard (http://broadinstitute.github.io/picard/; Version: 2.8.2) and SAMtools (Li et al., 2009) with default parameters.

To accurately identify SNPs, the low-quality and short alignments (mapping quality score <30 and length <30 bp) were filtered and BAM files were then indexed using SAMtools. SNP detection was performed using GATK HaplotypeCaller (DePristo et al., 2011) in -ERC GVCF mode, which finds and realigns indels during variant calling. Then, all genomic VCFs (gVCFs) were combined for joint genotyping with CombineGVCFs which produces a set of joint-called SNP and indels ready for filtering. To reduce the number of false positives, a high SNP confidence score was set with the following parameters: -stand_call_conf 30 -stand_emit_conf 40. We then split variants into SNPs and indels with SelectVariants and chose only biallelic SNPs with --restrictAllelesTo BIALLELIC. To ensure high quality variant calling, we used the following hard filters with VariantFiltration: QUAL < 30.0, SQR > 3.0, FS > 60.0 and MQ < 40.0, to obtain a single VCF file with unphased loci for the eight eastern African accessions.

We then used VCFtools (Danecek et al., 2011) to filter a published VCF file with 1217 accessions (Durvasula et al., 2017) and obtain only biallelic SNPs in the five nuclear chromosomes. We then added the 16 newly sequenced eastern African accessions to this VCF with BCFtools. After obtaining a merged VCF file with all accessions, we used Beagle (May 2020 version release) (Browning and Browning, 2009) to phase genotypes and impute missing SNPs based on linkage disequilibrium with default parameters. We used this VCF to produce a GDS file in the SNPRelate R package (Zheng et al., 2012). With this genofile, we obtained a list of polymorphic SNPs and a SNP matrix from the ecotypes here studied. These files were used to produce a BED file from the list of SNPs with a custom R code, and to assess Identity by State and produce an ibs matrix for all loci to account for population genetic structure in GWAS.

**S2. Growing medium in large scale experiments (1–3).** In Experiment 1 and 2 pots were filled with 50% autoclaved growing media (PGX PRO-MIX, Premier Tech Manufacturer), 25% commercial grade sand, and 25% calcined clay, with 2.5 cm of potting mix as topsoil to improve germination and initial growth. In Experiment 3, proportions were 20% autoclaved PGX, 40% sand, and 40% calcined clay. We covered pots with Press'n Seal Wrap to prevent dehydration, placed them under experimental conditions below, then removed covers after germination.

### Supplemental Results

**S1. Analysis of RGB color data.** To characterize the multidimensional color variation of leaves (or their extracts) among genotypes and environments in Experiments 2 and 3, we calculated principal components of RGB color (Fig. 2 B and C), revealing distinct patterns between experiments. In Experiment 2, higher amounts of red + green light loaded positively on $PC1_{LeavesColor}$ while higher amounts of blue light loaded positively on $PC2_{LeavesColor}$. Thus, under high light leaves were darker (lower values of PC1) and bluer (higher values of PC2) than under low light where leaves were light green, probably due to reduced pigments (Fig. 2 B). For leaf extracts in Experiment 3, higher amounts of blue + green light loaded positively on $PC1_{ExtractsColor}$ while red light loaded negatively on $PC2_{ExtractsColor}$. Thus, under night-time freezing extracts were less green (lower values of PC1) and dark orange to deep red (lower values of PC2), suggesting more anthocyanins and carotenoids than chlorophylls, while under no freezing extracts tended to be light green/orange, likely indicative of chlorophylls and xanthophylls (Fig. 2 C). Photographs taken on rosettes prior to harvest in both experiments showed that plants were darker/bluer under stress from high light and night-time freezing, thus extracts from leaves probably give greater information on pigment diversity than RGB color assessments from leaf photographs.

**S2. Flowering Time GWAS in northwestern Europe ecotypes**. In northwestern Europe, the top QTL for flowering time at 10 and 16ºC tagged a region expanding ca. 13.4 kb including *FLOWERING LOCUS C* (*FLC*), its upstream region (*SOK2*), and the enzyme *myo-inositol-1-phosphate synthase 3* (*MPSI3*). *FLC* is a flowering repressor whose function is downregulated via chromatin changes induced by the cold-responsive *FRI* or *VIN3*, two pathways that repress *FLC* during vernalization (Kim and Sung, 2013). *MPSI3* coregulates with *MPS1* and *MPS2* myo-inositol homeostasis and has roles in growth and development, with loss-of-function mutants having reduced size and delayed flowering under cold conditions (Fleet et al., 2018). The 2$^{nd}$ top QTL for flowering time at 10ºC was absent from the top 100 SNPs associated with flowering time at 16ºC. This QTL extended a region of ca. 466 kb including *VIN3* and flanking regions that encompassed *SUPPRESSOR OF MORE AXILLARY GROWTH2 5* (*SMXL5*) in one end, and *SUPPRESSOR OF MAX2 1* (*SMAX1*) in the other end, proteins that regulate various physiological processes involved in seed dormancy response to cold, growth and development (Stanga et al., 2013; Temmerman et al., 2022). The 3$^{rd}$ top QTL (flowering time at 10ºC) and 2$^{nd}$ top QTL (flowering time at 16ºC) tagged *SHORT VEGETATIVE PHASE* (*SVP*), a flowering repressor that functions within the thermosensory pathway in conjunction with *FLOWERING LOCUS M* (Jin et al., 2022). On the other hand, in central Europe/Caucasus, the 5$^{th}$ and 6$^{th}$ top QTL for flowering time at 10 and 16ºC, respectively, were upstream of the enzyme *myo-inositol-1-phosphate synthase 3* (*MPSI3*).

**S3. Physiological effects related with antioxidants under high light of *PHT4;4* variants**. Using ANOVA, we found strong treatment effects (high-light vs. low-light, under cold) on antioxidant activity, which was higher in all genotypes under high vs. low light grown in cold ($p$<0.0002 for knockout contrasts and $p$=0.01 for ecotypes contrast) (Supplemental Figure S8 A).

Chlorophyll content showed strong GxE in CS444342 and Col-0 (*p*=0.01), and Salk_082875 and Col-0 followed a similar trend (*p*=0.2): chlorophyll in the wild-type did not change in response to high light, but chlorophyll was reduced under high light in insertion lines, while it was even lower in the western Mediterranean ecotypes (Supplemental Figure S8 B).

We summarized leaf extracts RGB colors with PCA and used the first and second PCs and RGB brightness as our color traits. PC1 explained 81.1% of the variation and was strongly negatively correlated with RGB brightness (i.e., transparency of extract color) under both light conditions (cor= –0.98, p<0.0001): lower values of PC1 corresponded to brighter/more transparent extracts. PC1 mostly separated western Mediterranean ecotypes under high light, regardless of their allele (more transparent extracts), from everything else. Also, under high light PC1 was positively correlated with chlorophyll content (cor=0.6, p<0.001), *i.e.*, more transparent extracts had lower chlorophyll content. PC1 showed strong GxE in CS26443 and Ler (*p*=0.01), where extracts' colors were more transparent (less chlorophyll) in the insertion line under high light, but not different in the wild-type, confirming *PHT4;4* effects on pigment response to high light (Miyaji et al., 2015) and cold as we show here. PC2 explained 15.7% of the variation and mostly separated the treatment effect: leaf extracts were green, yellow, to orange (high values of PC2) and under low light leaf extracts were dark green, blue to gray (low values of PC2). PC2 was strongly correlated with antioxidant activity (cor=0.63, p<0.0001): higher values of PC2 corresponded to extracts with higher antioxidant activity under high light.

### Supplemental Figures

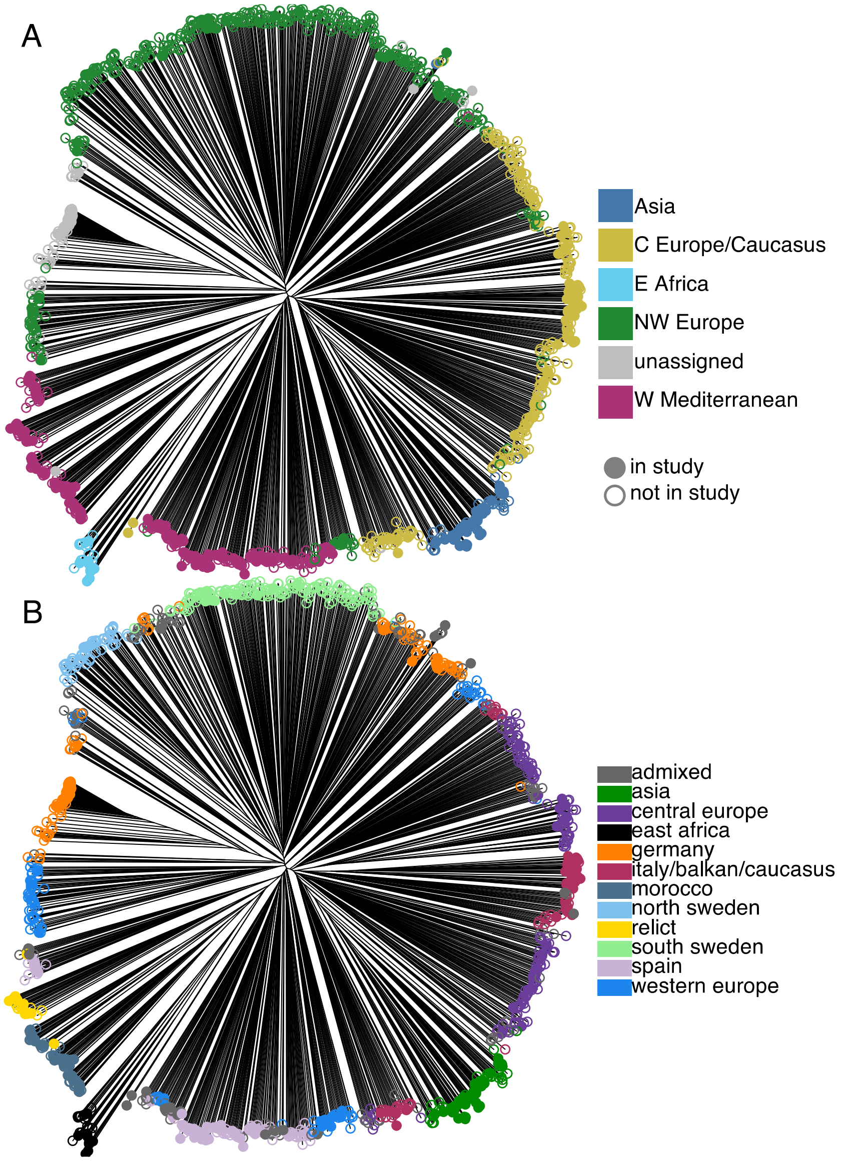


**Supplemental Figure S1** Neighbor-joining tree of 1233 Arabidopsis ecotypes, including 8 newly sequenced from eastern Africa, based on 540,351 resequencing SNPs filtered for LD and maf<0.05. Colors indicate: **A** regional classification used in this study (unassigned ecotypes are outside of the native range) and **B** genetic clusters previously circumscribed in Arabidopsis (Alonso-Blanco, 2016; Fulgione and Hancock, 2018). Filled circles are the 261 sequenced ecotypes used in this study. Data for this figure supports Figure 1 A.

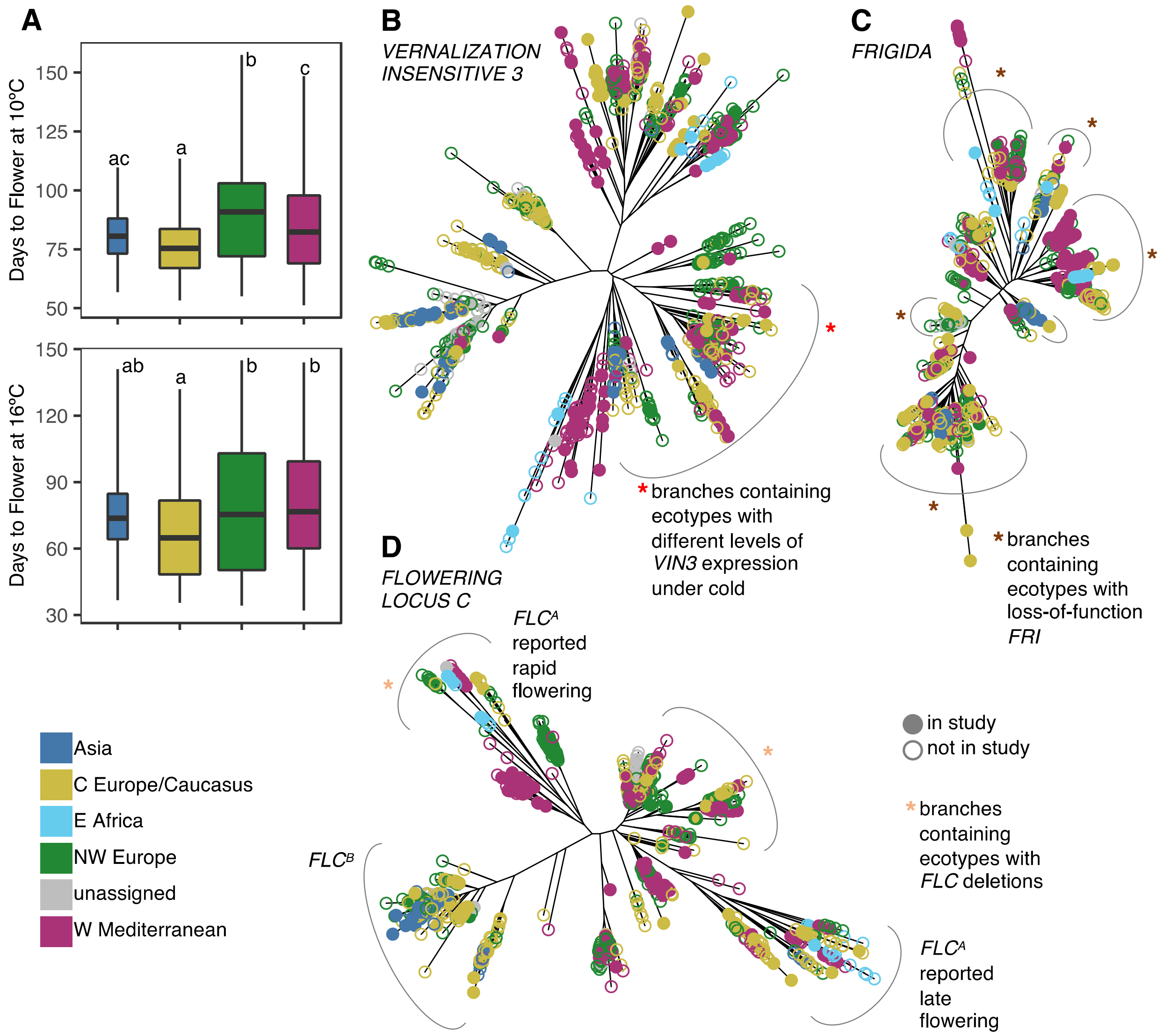


**Supplemental Figure S2** Flowering time variation based on 910–943 ecotypes and neighbor-joining trees of vernalization pathway genes based on 1233 Arabidopsis ecotypes and the gene coordinates ± 2.5 kbp. **A** Days to flower in plants grown at 10ºC (top) and 16ºC (bottom). Flowering data is based on Alonso-Blanco et al. (2016). Boxplots are proportional to sample size and depict the median and interquartile range, whiskers cover the data extent. **B** *VIN3*: AT5G57380, 819 SNPs in chr 5: 23243895–23252004 bp. **C** *FRI*: AT4G00650 1090 SNPs in chr 4: 266401–274003 bp. **D** *FLC*: AT5G10140, 1470 SNPs in 3170882–3181948 bp. This figure supports data for Figure 1 A. *VIN3* and/or *FRI* on *FLC* expression, which can also be affected with cold exposure alone. The data supporting branches marked with a red asterisk and the haplotype names for *FLC* comes from a literature review of relevant studies reporting natural genetic variation in these genes (see References for Supplemental Figure S2). Data for this figure supports Figure 2 A.

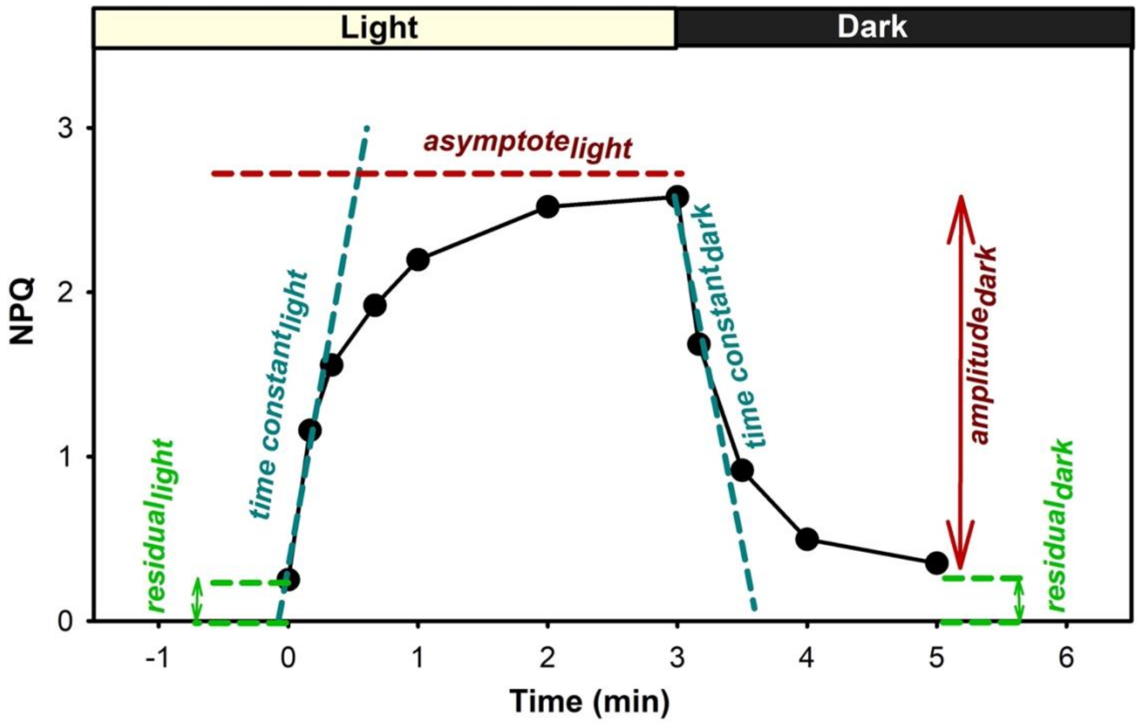


**Supplemental Figure S3** Representative NPQ induction and relaxation curve showing NPQ kinetics parameters. By fitting the NPQ data to exponential equations, six distinct parameters were obtained: ***residual**$_{light}$* - initial NPQ prior to light exposure; ***time constant**$_{light}$* - time needed to achieve 63% of maximum capacity of NPQ induction following light exposure; ***asymptote**$_{light}$* - maximum value of NPQ estimated from the asymptote; and when light was removed ***time constant**$_{dark}$* - time needed to achieve 63% of NPQ relaxation in dark; ***amplitude**$_{dark}$* - range of NPQ response in dark and ***residual**$_{dark}$* - not relaxed NPQ at the end of the dark. This figure supports data for Figure 3 and 8 and S4 and S5.

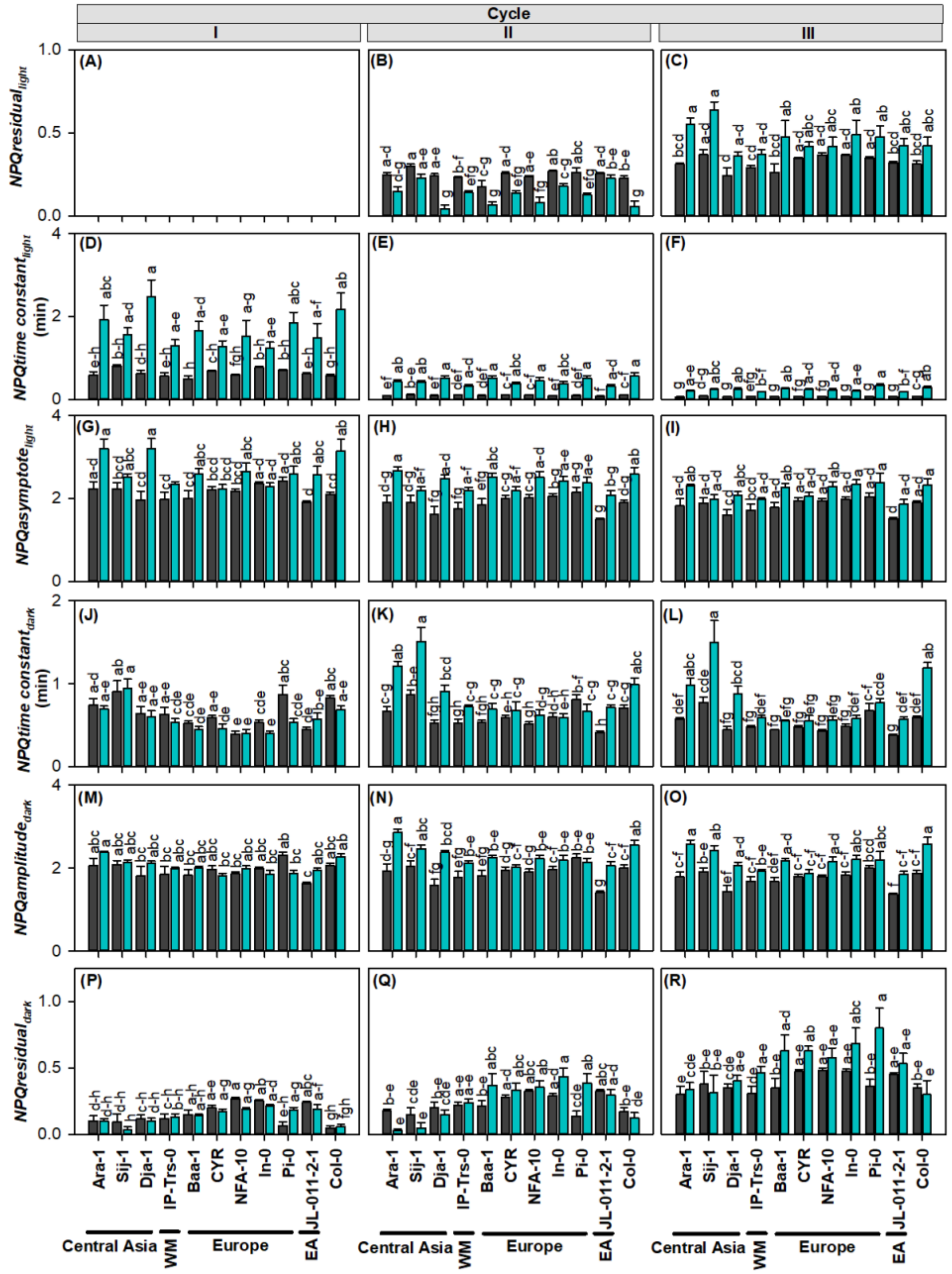


**Supplemental Figure S4** Effect of cold on NPQ kinetics for three subsequent light-dark cycles in 11 ecotypes. (A–C) *residual*$_{light}$ - the initial NPQ prior to light exposure; (D–F) *time constant*$_{light}$ - time needed to achieve 63% of maximum capacity of NPQ induction following light exposure; (G–I) *asymptote*$_{ligh}$ - maximum value of NPQ estimated from asymptote; and then when light was removed (J–L) *time constant*$_{dark}$ - time needed to achieve 63% of NPQ relaxation in dark (M–O) *amplitude*$_{dark}$ - range of NPQ response in dark; and (P–R) *residual*$_{dark}$ - not relaxed NPQ at the end of the dark, , with each estimated for three sequential light-dark cycles. Cyan bars represent cold treatment and gray bars represent control. Error bars indicate SEM (N=5–6 replicates). Different letters indicate significant differences from a post-hoc Tukey HSD test ($p < 0.05$). WM: western Mediterranean; EA: eastern Africa. This figure supports data for Figure 3 (left panels).

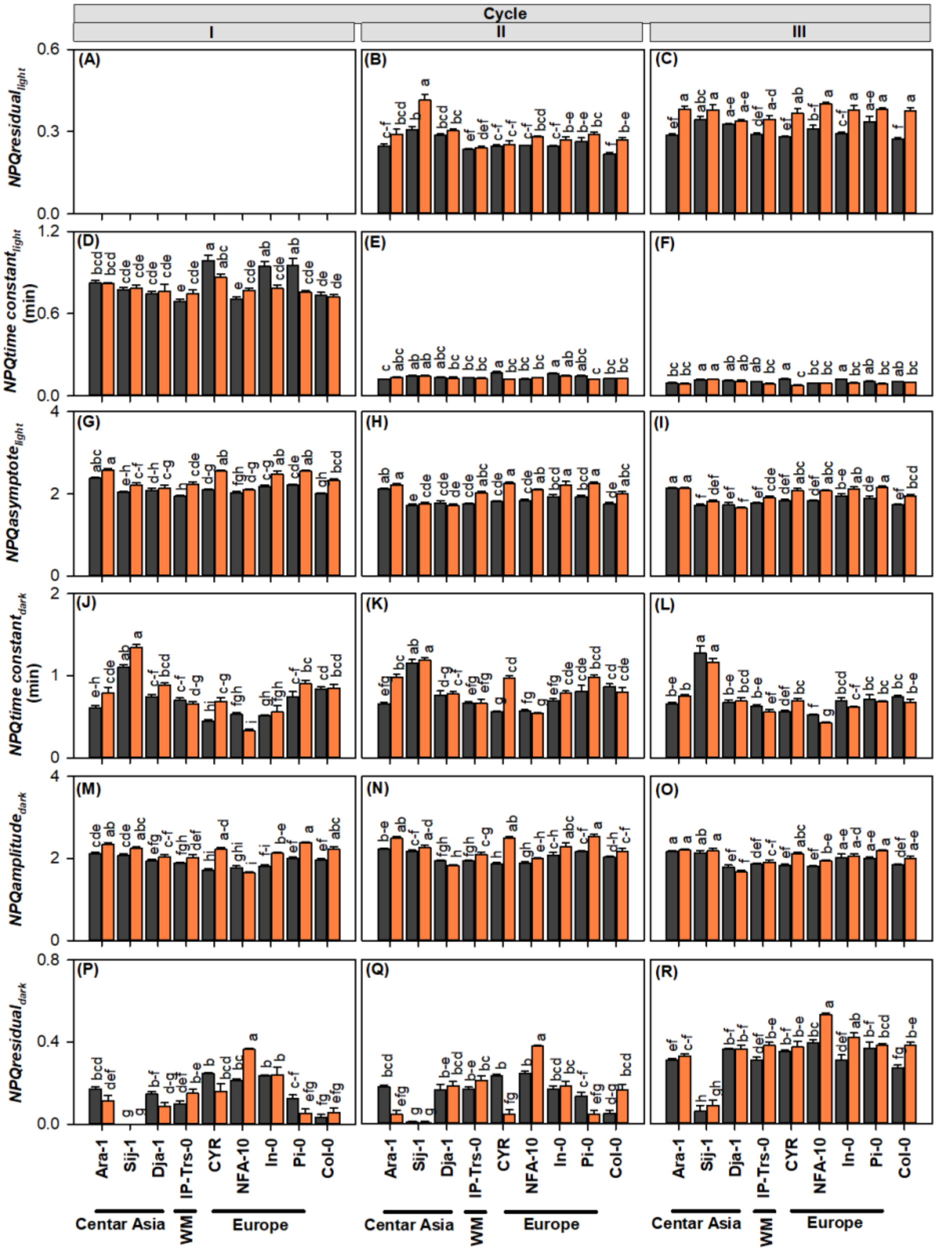


**Supplemental Figure S5** Effect of drought on NPQ kinetics for three subsequent light-dark cycles in 9 ecotypes (2 of 11 excluded due to flowering). (A–C) $residual_{light}$ - the initial NPQ prior to light exposure; (D–F) *time constant*$_{light}$ - time needed to achieve 63% of maximum capacity of NPQ induction following light exposure; (G–I) *asymptote*$_{ligh}$ - maximum value of NPQ estimated from asymptote; and then when light was removed (J–L) *time constant*$_{dark}$ - - the time needed to achieve 63% of NPQ relaxation in dark (M–O) *amplitude*$_{dark}$ - range of NPQ response in dark ; and (P–R) *residual*$_{dark}$ - not relaxed NPQ at the end of the dark,, with each estimated for three sequential light-dark cycles. Orange bars represent drought and gray bars control. Error bars indicate SEM (N=8–10 replicates, 5 in Dja-1). Different letters indicate significant differences from a post-hoc Tukey HSD test ($p < 0.05$). WM: western Mediterranean. This figure supports data for Figure 3 (right panels).

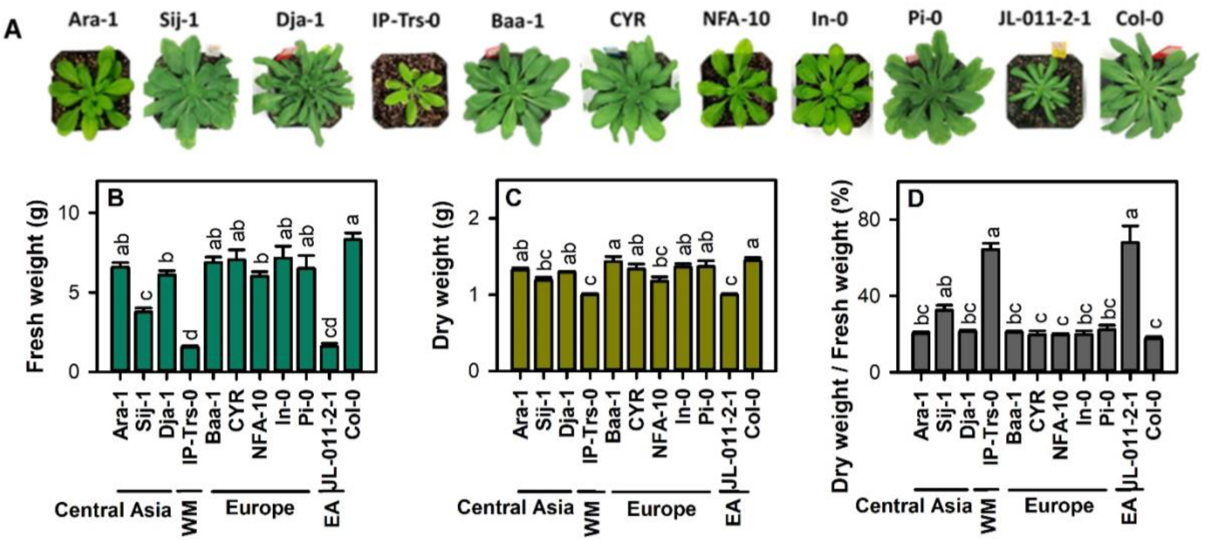


**Supplemental Figure S6** Aboveground biomass of 11 ecotypes: Ara-1 (~3000 m, but exact origin unclear), Sij-1 (2459 m), Dja-1 (2995 m), IP-Trs-0 (254 m), Baa-1 (22 m), CYR (52 m), NFA-10 (67 m), In-0 (1576 m), Pi-0 (2570 m), JL-011-2-1 (4078 m), and Col-0 (173 m); grown under control conditions and used in the NPQ kinetics under drought and cold experiments. **A** Representative picture of 21 d old plants. **B** Fresh weight. **C** Dry weight. **D** Dry matter content (%). Bars indicate mean and whiskers standard errors (N = 4–6 replicates). Different letters indicate significant differences from a post-hoc Tukey HSD test ($p < 0.05$). We indicate geographic origin of ecotypes. WM: western Mediterranean; EA: eastern African. This figure supports data for Figure 3.

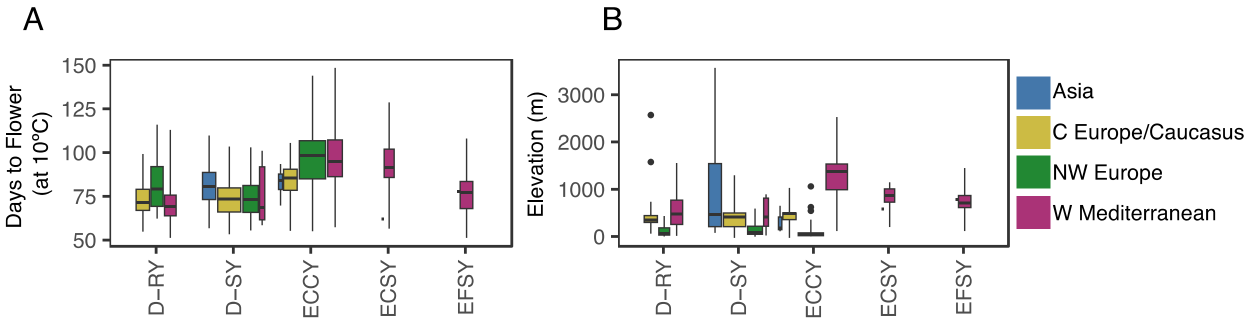


**Supplemental Figure S7** *DOG1* haplotype variation relative to flowering time (left) and elevation(right) in ecotypes from study regions. Boxplot width is proportional to haplotype frequency and depicts the median and interquartile range, whiskers extend to the data extremes. This figure supports data for Figures 5–7 and it is based on DOG1 published haplotype data (Martínez-Berdeja et al., 2020).

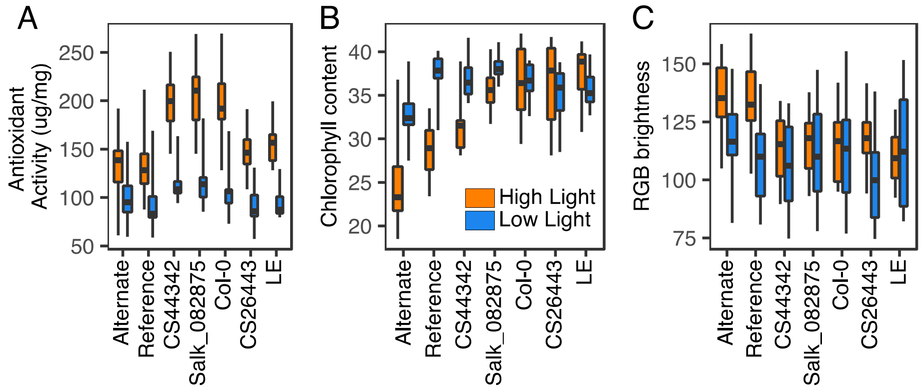


**Supplemental Figure S8** Physiological effects of *PHT4;4* variation in 20 d old plants exposed for 7 d to cold conditions (at 4ºC day and night) under high light (600 µmol $m^{-2}$ $s^{-1}$) vs. low light (150 µmol $m^{-2}$ $s^{-1}$). **A** Antioxidant activity (ug/leaf mg). **B** Chlorophyll content corresponding to relative chlorophyll content from SPAD. **C** Brightness based on red, blue, and green (RGB) values from photographs of leaf extracts. N ecotypes in the low antioxidant (alternate) GWAS allele = 3, N ecotypes in the high antioxidant allele = 2. Boxplot width is proportional to haplotype frequency (N=10–16 biological replicates per genotype) and depict the median and interquartile range, whiskers extend to the data extremes. This figure supports data for Figure 8.

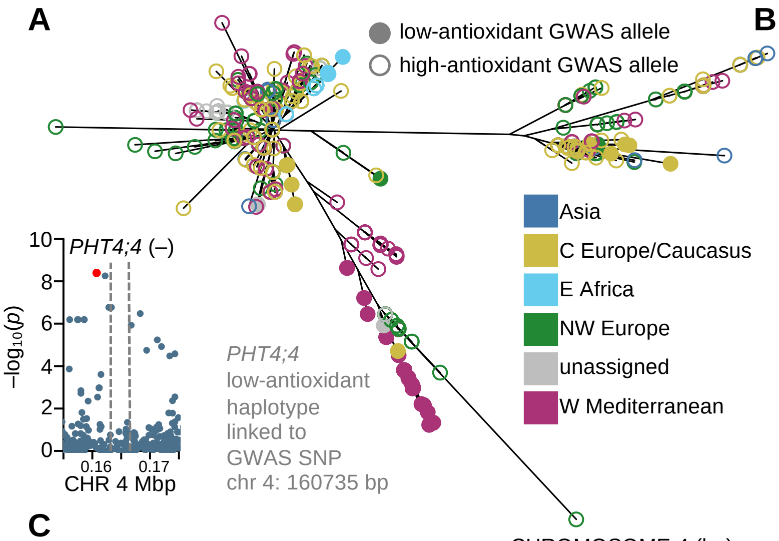


**B**

| MotifID | sequence | TFBS family |
|---|---|---|
| 1 | TCTACATTAATGAGG | HD-ZIP |
| 3 | AAAGACATAATTAAG | DOF |
| 8 | AAAAAAGCAAGGT | DOF |
| 11 | ATTAACTAAAAATGA | DOF |
| 12 | TAAAAGAGAGAGAGA | DOF |
| 16 | GTAAAAAAAATGATT | DOF |
| 17 | AAAAACAACAACATA | DOF |
| 19 | AAAACAAAGTCACTA | DOF |
| 24 | ACAGAGAATAGAAGA | DOF |
| 32 | ATAGTATAAAATAGG | DOF |
| 37 | CCAAAAAAAAACTTA | DOF |
| 40 | CTTTTCCATTTTAGT | DOF |
| 45 | TCTACATTTTTTTGT | DOF |
| 47 | TACAATAACAAAG | DOF |

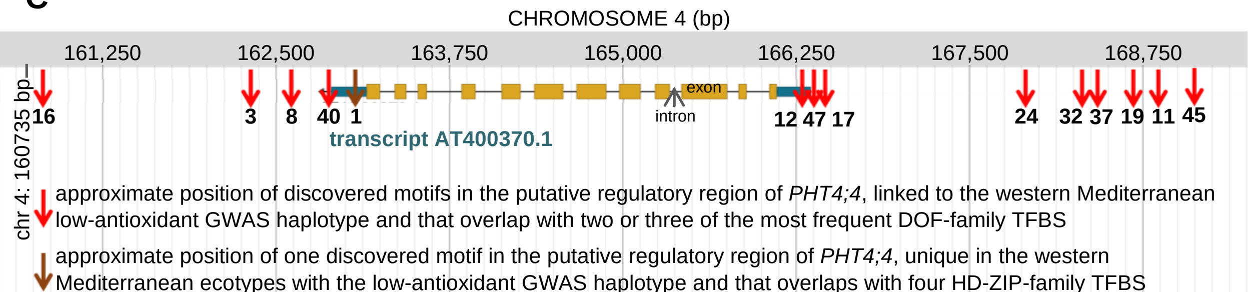


**Supplemental Figure S9** *PHT4;4* low-antioxidant GWAS haplotype and associated motifs with known transcription factors binding sites (TFBS). **A** Neighbor-joining tree of 1233 ecotypes using 251 SNPs from chr 4: 160735 bp (the 2nd top GWAS SNP for antioxidant activity under high light) to chr 4: 163308 bp (a *PHT4;4* intron SNP that was also a strong GWAS hit). 63 ecotypes with filled circles were selected from the branch inside the grey box, and contrasted with closely related ecotypes using the tree in Fig. S1, to discover *PHT4;4* putative *cis*-regulatory motifs associated with chr 4: 160735 bp. Zoomed-in manhattan plot shows with a box the haplotype genomic coordinates, while dashed lines delimits the *PHT4;4* coding region. Only ecotypes in the boxed branch were phenotyped in our high light GWAS experiment. **B** Selected discovered motifs in the putative *PHT4;4* regulatory region, indicating the TFBS family of significantly overlapping TFBS. **C** Graphical representation of the *PHT4;4* coding and regulatory regions with positions of motifs in B. This figure supports data for Table 3.

### Supplemental Tables

**Supplemental Table S1** Results of ANOVA for an effect of ecotype (E), cold or drought treatment (T), and genotype by environment interaction on 17 parameters of NPQ kinetics obtained by fitting an exponential equation to curves of NPQ induction (light) and relaxation (dark) in three subsequent light-dark cycles. The *p*-values are bolded for parameters where a significant effect of factor and/or factor interaction was detected. Two ecotypes were excluded from the drought experiment due to poor germination and early flowering.

| **Parameter** | ***p*-value** | | | | | |
|---|---|---|---|---|---|---|
| | Cold (11 ecotypes) | | | Drought (9 ecotypes) | | |
| | Ecotype (E) | Treatment (T) | E × T | E | T | E × T |
| *Light-dark Cycle I* | | | | | | |
| *NPQtime constant*$_{light}$ | **0.02** | **<0.0001** | **0.03** | **<0.0001** | **0.04** | **<0.0001** |
| *NPQasymptote*$_{light}$ | **0.01** | **<0.0001** | **0.001** | **<0.0001** | **<0.0001** | **0.002** |
| *NPQtime constant*$_{dark}$ | **<0.0001** | **0.007** | 0.08 | **<0.0001** | **0.002** | **<0.0001** |
| *NPQamplitude*$_{dark}$ | **<0.0001** | 0.1 | **0.003** | **<0.0001** | **<0.0001** | **<0.0001** |
| *NPQresidual*$_{dark}$ | **<0.0001** | 0.3 | **0.04** | **<0.0001** | 0.6 | **<0.0001** |
| *Light-dark Cycle II* | | | | | | |
| *NPQresidual*$_{light}$ | **<0.0001** | **<0.0001** | 0.08 | **<0.0001** | **<0.0001** | **0.01** |
| *NPQtime constant*$_{light}$ | **0.0003** | **<0.0001** | **0.04** | **<0.0001** | **0.008** | **<0.0001** |
| *NPQasymptote*$_{light}$ | **0.0003** | **<0.0001** | 0.05 | **<0.0001** | **<0.0001** | **0.0009** |
| *NPQtime constant*$_{dark}$ | **<0.0001** | **<0.0001** | **<0.0001** | **<0.0001** | **<0.0001** | **<0.0001** |
| *NPQamplitude*$_{dark}$ | **<0.0001** | **<0.0001** | **<0.0001** | **<0.0001** | **<0.0001** | **<0.0001** |
| *NPQresidual*$_{dark}$ | **<0.0001** | 0.1 | **0.0002** | **<0.0001** | 0.05 | **<0.0001** |
| *Light-dark Cycle III* | | | | | | |
| *NPQresidual*$_{light}$ | **0.002** | **<0.0001** | 0.2 | **<0.0001** | **<0.0001** | **0.0006** |
| *NPQtime constant*$_{light}$ | **0.006** | **<0.0001** | **0.01** | **<0.0001** | **<0.0001** | **<0.0001** |
| *NPQasymptote*$_{light}$ | **<0.0001** | **<0.0001** | 0.6 | **<0.0001** | **<0.0001** | **0.0001** |
| *NPQtime constant*$_{dark}$ | **<0.0001** | **<0.0001** | **0.0002** | **<0.0001** | 0.05 | **<0.0001** |
| *NPQamplitude*$_{dark}$ | **<0.0001** | **<0.0001** | **0.02** | **<0.0001** | **<0.0001** | **0.006** |
| *NPQresidual*$_{dark}$ | **<0.0001** | **<0.0001** | 0.07 | **<0.0001** | **<0.0001** | **0.002** |

**Supplemental Table S2** Quantification of photosynthesis related pigments in 8 ecotypes (3 of 11 excluded due to flowering). Numbers represent means and SEM are indicated in brackets (*N* = 4–6 biological replicates). The *p*-values indicate the effect of the ecotype. Different letters indicate significant differences from a post-hoc Tukey HSD test ($p \leq 0.05$). Car - total carotenoids; chl *a* - chlorophyll *a*; chl *b* – chlorophyll *b*.

| | **Ecotype** | | | | | | | | **P-value** |
|---|---|---|---|---|---|---|---|---|---|
| | **Central Asia** | | **Europe** | | | | **East Africa** | | |
| | **Sij-1 (2459 m)** | **Dja-1 (2995 m)** | **Baa-1 (22 m)** | **CYR (52 m)** | **In-0 (1576 m)** | **Pi-0 (2570 m)** | **JL11-2-1 (4078 m)** | **Col-0 (173 m)** | |
| Chl *a* ($g\ m^{-2}$) | $0.250^{a}$ (0.014) | $0.217^{ab}$ (0.005) | $0.230^{ab}$ (0.008) | $0.199^{bc}$ (0.004) | $0.209^{bc}$ (0.009) | $0.229^{ab}$ (0.007) | $0.128^{c}$ (0.003) | $0.229^{ab}$ (0.010) | 0.0007 |
| Chl *b* ($g\ m^{-2}$) | $0.124^{a}$ (0.007) | $0.108^{ab}$ (0.009) | $0.108^{ab}$ (0.004) | $0.088^{bc}$ (0.004) | $0.098^{abc}$ (0.011) | $0.110^{ab}$ (0.002) | $0.067^{c}$ (0.001) | $0.103^{abc}$ (0.003) | 0.0009 |
| Chl *a* + *b* | $0.374^{a}$ (0.021) | $0.325^{ab}$ (0.013) | $0.338^{ab}$ (0.010) | $0.287^{bc}$ (0.006) | $0.307^{bc}$ (0.019) | $0.339^{ab}$ (0.008) | $0.195^{c}$ (0.004) | $0.333^{bc}$ (0.012) | 0.0007 |
| Chl *a* / *b* | 2.03 (0.05) | 2.06 (0.16) | 2.14 (0.07) | 2.28 (0.10) | 2.19 (0.19) | 2.08 (0.04) | 1.92 (0.02) | 2.22 (0.08) | 0.05 |
| Total Car ($g\ m^{-2}$) | $0.0532^{a}$ (0.0014) | $0.0353^{cd}$ (0.0012) | $0.0412^{bc}$ (0.0015) | $0.0347^{d}$ (0.0011) | $0.0350^{cd}$ (0.0007) | $0.0398b^{cd}$ (0.0012) | $0.0202^{e}$ (0.0006) | $0.0423^{b}$ (0.0022) | <0.0001 |
| Chl *a* + *b* / Total Car | $7.04^{c}$ (0.40) | $9.26^{ab}$ (0.56) | $8.22^{abc}$ (0.26) | $8.32^{cb}$ (0.29) | $8.76^{ab}$ (0.45) | $8.52^{abc}$ (0.14) | $9.65^{a}$ (0.17) | $7.91^{bc}$ (0.22) | 0.002 |

**Supplemental Table S3** Names and sequences of primers used for PCR to verify homozygosity of Arabidopsis insertional mutants in *PHT4;4*. CS26443 is a *Ds* transposon insertional mutant in a *Landberg erecta* (LE) background. Salk_082875 and CS444342 are T-DNA insertional mutants in a Col-0 background.

| **Mutant** | **Reaction product** | **Left primer** | **Right primer** |
|---|---|---|---|
| CS26443 | Wild-type (LE) allele | PHT4-L2 5′-ATGGAGATGCGTTCTGTAGATT-3′ | PHT4-R 5′-GGTTCCAACGAGTAGAAGATGA-3′ |
| | Mutated allele (targeting *Ds* transposon) | Ds3-4 5'- CCGTCCCGCAAGTTAAATATG -3' | PHT4-L2 or PHT4-R |
| Salk_082875 | Wild-type (Col-0) allele | Salk_082875_LP 5′-AGTCAACACACCCATTGAAGC -3′ | Salk_082875_RP 5′- ATGTGGTGAAGTCGACCGTAG -3′ |
| | Mutated allele (targeting T-DNA) | LBb1.3 5′- ATTTTGCCGATTTCGGAAC -3′ | Salk_082875_RP 5′- ATGTGGTGAAGTCGACCGTAG -3′ |
| CS444342 | Wild-type (Col-0) allele | GABI_46H02_LP 5′- GGTAGAAGGGGACCCTTGAAC -3′ | GABI_46H02_RP 5′- CGAACTTCTTGGACATCTTCG -3′ |
| | Mutated allele (targeting T-DNA) | Sul7 5′- GACTACAGTCAGCCGTGCTTC -3′ | Sul9 5′- GGTTTCCGAGATGGTGATTG -3′ |
